# Efficient CRISPR/Cas9-mediated homology independent sequence replacement in vivo and non-dividing primary cells

**DOI:** 10.64898/2026.07.24.740048

**Authors:** Tu Dang, Alexandra Roman, Anja Zimmer, Mikhail Lebedin, Ella Bahry, Claudia Marti Grifol, Markus Esser, Seren Sevim Wunderlich, Duncan C. Miller, Sebastian Diecke, Ralf Kühn

## Abstract

Precise sequence replacement in non-dividing cells remains a major challenge for genome editing. Here we show that REPLACE (Rational end-joining protocol delivering a targeted sequence exchange), a homology-independent CRISPR/Cas9-based replacement strategy, enables exon- and gene-scale substitution in primary cells, *in vivo* tissues and post-mitotic human cardiomyocytes. REPLACE uses two guide RNAs to excise a defined genomic region and inserts a donor lacking homology arms through non-homologous end joining (NHEJ). In primary mouse hepatocytes, REPLACE mediated exon replacement in 35% of all cells. In adult mouse liver, editing efficiency could be increased to ∼20% when Cas9-sgRNA ribonucleoproteins were delivered via engineered virus-like particles (eVLPs) together with an adeno-associated virus (AAV) donor. REPLACE also supported large-segment replacement, enabling one-step exchange of a ∼27-kb mouse *Ace2* interval with the human *ACE2* coding region in zygotes, followed by germline transmission and tissue-specific expression. Finally, we applied REPLACE to a disease-relevant mutation that is not readily addressable by base editing and was poorly corrected by prime editing in post-mitotic cardiomyocytes. At the *LMNA* locus, REPLACE corrected the K117fs frameshift mutation in patient-derived post-mitotic cardiomyocytes with precise exon replacement and restored Lamin A/C protein expression and nuclear lamina localization. These findings establish REPLACE as a versatile platform for homology-independent sequence replacement and as a complementary approach for genetic correction in settings where homology-directed repair (HDR), base editing (BE) or prime editing (PE) are inefficient or not applicable.

## Introduction

The CRISPR/Cas9 system is a powerful tool for precise genome editing in mammalian cells. In this approach, single-guide RNAs (sgRNAs) direct Cas9 nucleases to a specific target sequence, allowing Cas9 nucleases to form double-strand breaks (DSBs) ^1,2^. Gene editing at Cas9 induced DSBs can be achieved by harnessing two primary cell repair pathways. Non-homologous end joining (NHEJ) typically introduces small insertions or deletions (indels), while homology-directed repair (HDR) uses an exogenous repair template to enable precise sequence modification ^3–5^. Because HDR is restricted to the S and G2 phases of the cell cycle and relies on the presence of a repair template, it occurs far less frequently than NHEJ and is greatly reduced or entirely absent in slowly or non-dividing cells ^6^. In contrast, NHEJ functions independently of the cell cycle phase, making it active in both proliferating and post-mitotic cells ^7^. This characteristic renders NHEJ a highly attractive tool for somatic gene therapies in tissue with limited regenerative capacity, such as the heart or brain, which are difficult to target by HDR-based technologies.

More recently, base editing and prime editing have expanded the genome-editing toolbox by enabling precise sequence changes without requiring DSBs or donor DNA ^8–10^. However, these approaches also have major constraints. Base editing is restricted to defined nucleotide conversions and is not suitable for arbitrary insertions, deletions, or exon-scale sequence replacement. Prime editing is generally more versatile, but its efficiency is highly context-dependent and can remain low in therapeutically relevant primary or post-mitotic cells ^11^.

Therapeutic strategies focusing on gene disruption by NHEJ are advancing in clinical trials, particularly for diseases where the deletion of pathogenic misfolded proteins can alleviate disease phenotypes. Such an approach was used to treat Transthyretin amyloidosis (ATTR), a life-threatening disease caused by a single nucleotide mutation in the *TTR* gene ^12^. Furthermore, NHEJ has been harnessed in several experimental and preclinical settings for targeted exon deletion or exon skipping in somatic muscle cells with the aim of treating Duchenne Muscular Dystrophy (DMD) ^13,14^. In 2023, the FDA approved the first CRISPR/Cas9-based treatment -Casgevy-for sickle cell disease (SCD) and transfusion dependent ß-thalassemia (TDT) ^14^. Casgevy’s strategy uses NHEJ to disrupt a transcriptional repressor of γ-globin in hematopoietic stem and progenitor cells (HSPCs), thereby reactivating fetal hemoglobin production prior to reimplantation of the modified cells ^15^.

However, the canonical NHEJ-based approaches for gene therapy focus largely on knockouts and exon skipping. While these strategies can be clinically relevant, they restrict the spectrum of possible edits. In 2016 Suzuki et al. developed an approach called Homology-independent targeted integration (HITI), which utilized the NHEJ repair pathway to mediate DNA integration in both dividing and non-dividing cells in vitro and in vivo ^16^. They demonstrated HITI’s ability for reporter tagging in postmitotic neurons as well as its potential for partially repairing genetic defects that result from large deletions.

Recognizing the robust, cell-cycle-independent activity of NHEJ yet aiming to overcome the inherent limitations of canonical NHEJ, we recently developed a novel NHEJ-based gene replacement approach called REPLACE (Rational end-joining protocol delivering a targeted sequence exchange) ^17^. REPLACE is a versatile and flexible strategy that enables a wide range of large gene modifications.

REPLACE exploits two sgRNAs that simultaneously guide the Cas9 nuclease to flank a genomic region of interest, thereby creating two double-strand breaks and excising the target sequence. Into this excised region, a donor sequence carrying the desired replacement, lacking homology arms, is then ligated via NHEJ. This approach bypasses the need for homology-directed repair, making it well-suited for post-mitotic or slowly dividing cells. The proof-of-concept for REPLACE was first reported in 2021 by Danner et al., demonstrating successful sequence replacement in various immortalized cell lines, including HEK293, HeLa, and K-562 cells ^17^.

Here, we expand upon the initial work of Danner et al. by showing, for the first time, that REPLACE can also mediate efficient exon replacement in primary, non-dividing cells. We recorded targeted exon exchange with up to 35% editing efficiency in primary mouse hepatocytes and with up to 20% in vivo editing in the adult liver of mice. Additionally, we demonstrate the feasibility of replacing large gene segments by exchanging a 27-kilobase (kb) gene interval of the mouse *Ace2* gene with the human homolog’s coding region. Finally, we applied REPLACE to the clinically relevant *LMNA* locus in patient-derived post-mitotic cardiomyocytes, correcting the K117fs frameshift mutation, which is not readily addressable by base editing and was poorly corrected by prime editing in this cellular context ^11^. REPLACE-mediated correction restored Lamin A/C protein expression and nuclear lamina localization. Collectively, these findings establish REPLACE as a versatile homology-independent sequence replacement platform and a complementary approach for genetic correction in settings where HDR or nicking-based approaches are futile or unsuitable.

## Results

### 1. REPLACE mediates exon replacement in primary cells

REPLACE editing aims to exchange a genomic sequence with a DNA double-stranded donor without relying on homology arms. In this approach, two sgRNAs simultaneously guide the Cas9 nuclease to sites flanking a genomic region of interest, generating two double-strand breaks (DSBs). The excised genomic region is then replaced with a donor cassette via NHEJ (Fig. 1a).

**Figure 1.**
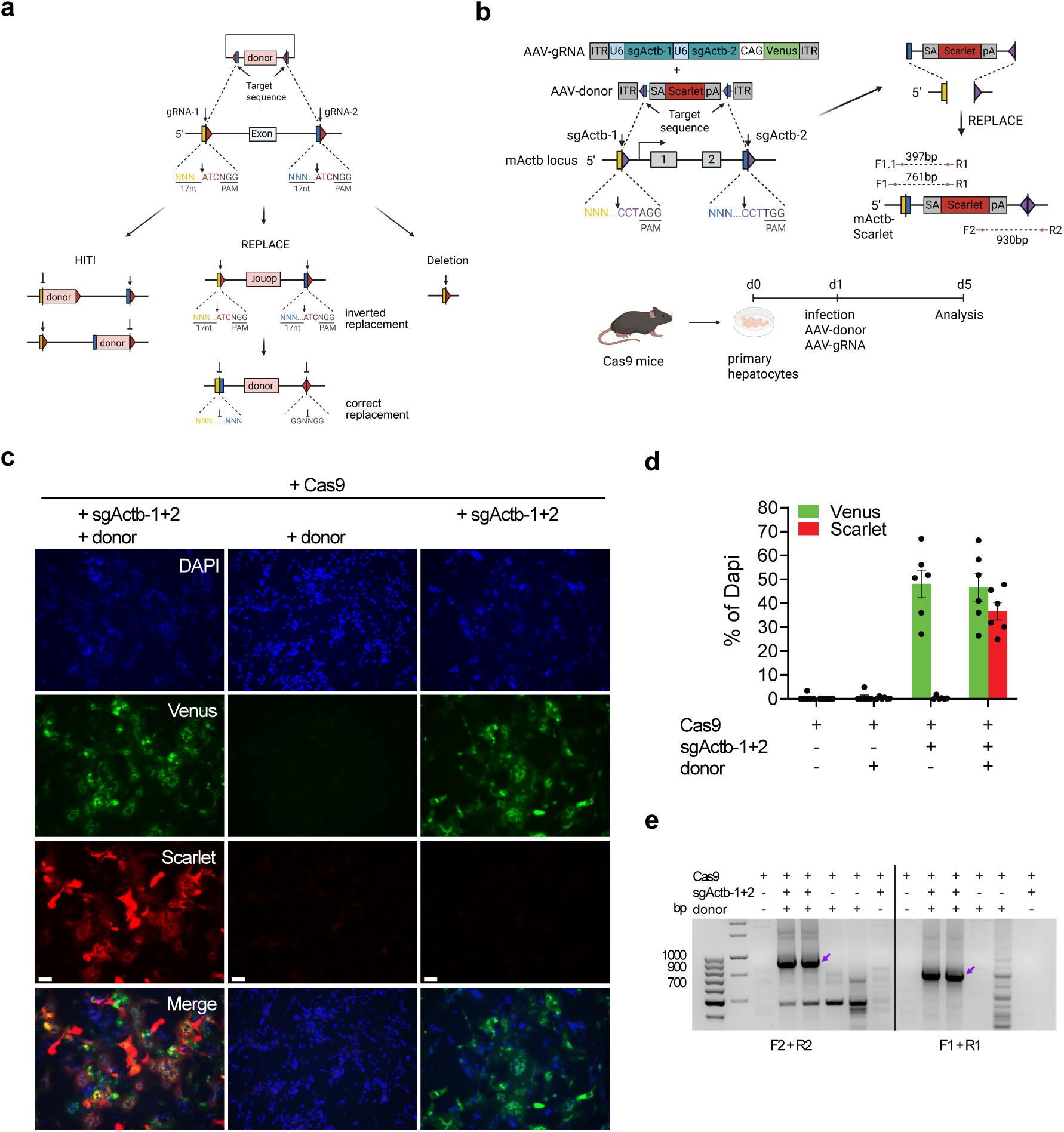
REPLACE enables efficient exon replacement in primary mouse hepatocytes. **(a)** Scheme illustrating the targeting concept of REPLACE. Pentagons represent Cas9/sgRNA target sites, black arrows indicate the Cas9 cleavage site. Correct donor integration (correct replacement) disrupts the target site and is retained, whereas undesired outcomes (deletion or inverted replacement) can reconstitute a gRNA target site and may be re-cleaved, thereby enriching for the intended product. **(b)** Targeting strategy and experimental scheme for replacing *Actb* exons 1 and 2 with a Scarlet reporter cassette. sgRNAs (sgActb-1 and sgActb-2) are indicated as black arrows. Genotyping primers are shown as purple arrows and labeled F1, F1.1, F2, R1 and R2. **(c)** Representative immunofluorescence images of primary hepatocytes 4 days after in vitro transduction with AAV-ACTB-donor and AAV-sgActb1+2. Scale bar, 100 µm. **(d)** Quantification of transduction and editing efficiencies. Venus^+^ cells (Venus^+^/DAPI^+^) indicate uptake of the gRNA vector; Scarlet^+^ cells (Scarlet^+^/DAPI^+^) indicate successful editing. Data are mean ± SD from six biological replicates (n = 6 independent experiments). **(e)** PCR genotyping of editing outcome using primers indicated in panel **b**. Purple arrow indicates REPLACE-specific bands.

The donor cassette is typically delivered using either a minicircle (MC) or AAV vector. For successful replacement, the donor cassette must be released from its delivery vector, which can be achieved by including one of the gRNA target sequences directly upstream and downstream of the cassette. Cleavage at these sites liberates the donor fragment for integration (Fig. 1a).

To further enhance REPLACE efficiency, we selected sgRNAs whose last three nucleotides (positions 18–20) are identical. This feature facilitates the reconstruction of one of the two original gRNA target sites in undesired products such as deletions or inverted donor replacement, allowing Cas9 to cleave these intermediates again and thereby favor the desired replacement event (Fig. 1a). Notably, reconstitution of the target site only occurs in a subset of editing outcomes, where re-ligation proceeds without introducing insertions or deletions at the DSB sites.

To evaluate REPLACE editing in non-dividing cells in vitro, we adapted the strategy from Danner et al. ^17^ and designed a cassette to replace two exons of the mouse *Actb* gene with a Scarlet-reporter (hereafter referred to as ACTB-Scarlet REPLACE).

First, we designed a pool of sgRNAs targeting the regions flanking exons 1 and 2 of the mouse *Actb* gene. A pair of sgRNAs is suitable for REPLACE if both guides show robust activity at their respective target sites. To identify effective sgRNA pairs, we transfected mouse neuroblastoma (N2A) cells with plasmids expressing SpCas9 and each candidate sgRNA pair. Three days post-transfection, we amplified the genomic region encompassing the two gRNA target sites to assess editing activity. In control (untreated or Cas9-only) and all treated samples, we detected an amplicon of 1128 bp, representing either unedited alleles or alleles cleaved at only one site. When both sgRNAs were active, we expected an additional shorter PCR fragment resulting from the excision of the intervening genomic region. Only the combination of sgRNA-1 and sgRNA-2 (here referred to as sgActb-1 and sgActb-2) yielded this predicted shorter band of ∼368 bp (Supplementary Fig. 1a), suggesting its suitability for subsequent ACTB-Scarlet REPLACE experiments.

Next, we tested a DNA donor cassette (ACTB-donor) containing the endogenous mouse ACTB splice acceptor, followed by the coding sequence for the fluorescent protein Scarlet and a polyadenylation (poly-A) signal (Fig. 1b). This cassette is flanked by the sgActb-2 target site in the reverse orientation. Because this orientation prevents the sgActb-2 site from being reconstituted in the intended editing outcome (correct replacement), we hypothesize it will reduce unproductive re-cleavage events, thereby enhancing REPLACE efficiency.

To evaluate REPLACE editing in primary non-dividing cells in vitro, we delivered the ACTB-donor and the sgRNAs sgActb-1 & 2 using two AAV-serotype D/J vectors (Fig. 1b). Both sgRNAs were cloned into a single AAV-D/J vector (hereafter referred to as AAV-sgActb1+2) under the control of the U6 promoter. To monitor the infection efficiency of the AAVs, we introduced a Venus reporter, driven by the CAG promoter, into the same vector. Next, we isolated hepatocytes from mice (R26-Cas9) constitutively expressing Cas9 and transduced these primary hepatocytes with the REPLACE constructs (AAV-ACTB-donor: MOI 500,000 and AAV-sgActb1+2: MOI 500,000) one day after isolation (Fig. 1b). Four days post-transduction, numerous hepatocytes co-expressing Venus and Scarlet were observed in samples treated with both AAV-ACTB-donor and AAV-sgActb1+2, whereas control samples receiving either AAV-ACTB-donor or AAV-sgActb1+2 lacked Scarlet fluorescence (Fig. 1c). Venus expression confirmed successful transduction with AAV-sgActb1+2, while Scarlet expression indicated integration of the ACTB-donor at the target locus. Based on reporter expression, approximately 50% of the total cell population (relative to DAPI) was successfully infected with AAV-sgActb1+2, and REPLACE editing occurred in 37% of the total cell population (Fig. 1d). Restricting the calculation to Venus-positive cells alone revealed an editing efficiency of ∼74%, highlighting the robust potential of REPLACE editing in primary non-dividing hepatocytes.

Scarlet integration at the target locus can arise from either a correct replacement event (i.e., exchanging the donor sequence with the endogenous locus in the proper orientation) or integration of the ACTB-donor at the sgActb-1 target site via HITI. To distinguish between these two events, we designed a PCR-based genotyping strategy. We amplified the 5′ and 3′ integration junctions using primer pairs in which one primer binds inside the Scarlet reporter and the other in the genomic locus. While both correct replacement and HITI yield a 761 bp amplicon at the 5′ junction, the 3′ junction product differs: correct replacement produces a ∼930 bp band, whereas HITI results in a ∼2000 bp fragment. In samples treated with both AAV-ACTB-donor and AAV-sgActb1+2, we detected the expected 761 bp product at the 5′ junction and only the 930 bp product at the 3′ junction, indicating that the Scarlet expression observed predominantly arose from the correct replacement event rather than HITI (Fig. 1e).

To further validate that the genotyping amplicons represented correct replacement events, we subjected PCR products from both junction sites to Sanger sequencing (Supplementary Fig. 1b). The results confirmed that both amplicons contained the precise junction between the Scarlet cassette and the genomic locus, as predicted for correct replacement. Thus, our results demonstrate that REPLACE is an efficient exon replacement strategy in primary, non-dividing hepatocytes.

### 2. REPLACE mediates in vivo genome editing in the liver

To investigate whether REPLACE editing can be achieved in vivo, we first used R26-Cas9 mice which express Cas9 constitutively. Ten-week-old male R26-Cas9 mice were injected via the tail vein with AAV-D/J particles (2 x 10^10^ vg/g mouse for each construct) encoding the ACTB-donor and sgActb-1+2 (Supplementary Fig. 2a).

Eight weeks after injection, we evaluated the distribution and uptake of the REPLACE constructs by measuring in vivo fluorescence in multiple tissues, including the heart, skeletal muscle, adipose tissue, and liver. Cas9 mice that received AAV-ACTB-donor plus AAV-sgActb1+2 exhibited robust Venus fluorescence in the liver and a faint signal in the heart, but very little to no signal in skeletal muscle or adipose tissue, suggesting that most of the AAV constructs accumulated in the liver (Supplementary Fig. 2b). We also observed equally strong Scarlet fluorescence in hepatic tissue, suggesting that REPLACE editing had successfully occurred there. No Venus or Scarlet fluorescence was detected in the liver of control animals (untreated, PBS-injected, or AAV-ACTB-donor-only), confirming that the observed fluorescence arises from functional delivery and editing in R26-Cas9 mice (Supplementary Fig. 2c). To verify these observations at the cellular level, we performed fluorescence imaging on liver cryosections from mice injected with AAV-ACTB-donor and AAV-sgActb1+2. In these sections, we observed cells co-expressing Venus and Scarlet, confirming that both the viral delivery and editing occurred within the liver tissue (Supplementary Fig. 2d). Consistent with this result, primary hepatocytes isolated from the same animals also showed Venus and Scarlet co-expression upon ex vivo imaging, further supporting the conclusion that REPLACE editing was successfully achieved in hepatocytes in vivo (Supplementary Fig. 3a).

Flow cytometry analysis revealed that 26% of hepatocytes isolated from mice treated with AAV-ACTB-donor and AAV-sgActb1+2 were Venus-positive, confirming successful viral transduction in this cell fraction (Supplementary Fig. 3b). Among all hepatocytes, ∼13% were Scarlet-positive, representing the absolute editing frequency. When the analysis was restricted to Venus-positive cells only, up to 50% of transduced hepatocytes were also Scarlet-positive, indicating efficient REPLACE-mediated editing within the successfully targeted cell population (Supplementary Fig. 3b).

Next, we applied the same REPLACE targeting strategy to wild-type (WT) C57BL/6 (B6) mice. Since these animals do not express Cas9, we needed to provide the nuclease along with the donor and gRNAs. For this purpose, we adopted a trans-splicing intein system, wherein two separate AAV vectors each deliver one half of the Cas9 protein (Cas9-N & Cas9-C). When both fragments are expressed in the same cell, they reconstitute a full-length Cas9 nuclease.

We incorporated both sgRNAs (sgActb-1 and sgActb-2) into the Cas9-C fragment-expressing vector, while subcloning the donor into the Cas9-N fragment-expressing vector. This strategy yielded two AAV vectors in total for REPLACE in WT mice: AAV-sgActb-1+2-Cas9-C and AAV-ACTB-donor-Cas9-N (Fig. 2a). We then performed the in vivo REPLACE experiment by injecting these two vectors into 10-week-old male B6 mice. At 18 weeks of age, the animals were sacrificed, and primary hepatocytes were isolated. Approximately 11% of the total hepatocytes obtained from mice that received both AAVs were Scarlet-positive, representing a significant difference compared to control samples (PBS-injected or control gRNA and donor) (Fig. 2c-d).

**Figure 2.**
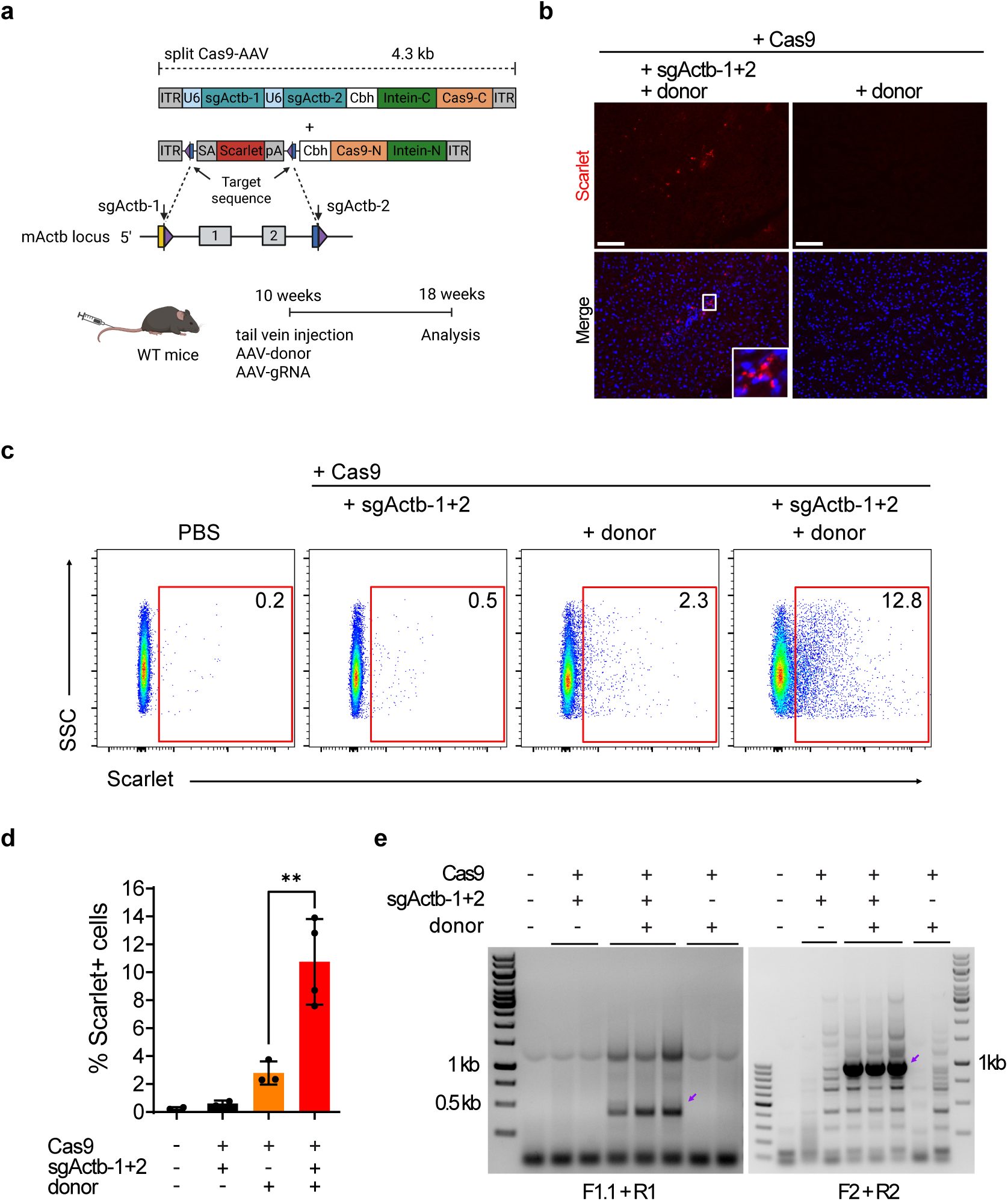
REPLACE mediates in vivo exon replacement in adult mouse liver. **(a)** Scheme illustrating the targeting concept of REPLACE at the *Actb* locus in adult mice using dual AAV-DJ vectors encoding splitintein Cas9. Both sgRNAs, sgActb-1 and sgActb-2, were encoded on the Cas9-C vector (AAV-sgActb1+2-Cas9-C) and the ACTB-donor cassette was encoded on the Cas9-N vector (AAV-ACTB-donor-Cas9-N). Ten-week-old male WT mice were injected via the tail vein with the indicated AAV combinations and subjected to downstream analysis 8 weeks post-injection. **(b)** Representative immunofluorescence images of adult liver tissue 8 weeks after injection with split-Cas9 AAVs. Scale bar, 100 µm. **(c–d)** Flow-cytometry analysis of Scarlet-expressing hepatocytes isolated from mice treated with the splitintein Cas9 REPLACE strategy compared with control groups (PBS-injected, Cas9+gRNAs and Cas9+donor). Gates indicate the percentages of Scarlet^+^ cells. Editing efficiency was measured as the percentage of Scarlet^+^ hepatocytes among total hepatocytes. Data are presented as mean ± SD (n = 2-4 mice per group). Statistical significance was evaluated using an unpaired two-tailed t-test with **P ≤ 0.01. **(e)** PCR genotyping of bulk edited hepatocytes using primers indicated in **Figure 1b**. Purple arrow indicates REPLACE-specific bands.

To distinguish between ACTB-donor integration (HITI) and correct exon replacement (REPLACE) among Scarlet^+^ cells, we used the same genotyping strategy described in Section 1. FACS-sorted Scarlet^+^ hepatocytes from animals treated with AAV-sgActb-1+2-Cas9-C and AAV-ACTB-donor-Cas9-N displayed the expected 397 bp PCR product at the 5′ junction - common to both HITI and REPLACE. However, only the 930 bp band appeared at the 3′ junction, which specifically indicates correct exon replacement (Fig. 2e). These data confirm that the observed Scarlet expression primarily arose from proper REPLACE events rather than HITI. Altogether, these results demonstrate the utility of REPLACE for exon replacement in post-mitotic hepatocytes in vivo.

### 3. Efficient in vivo replacement mediated by REPLACE using engineered virus-like-particles

In our previous experiments (Section 2), we demonstrated that REPLACE editing could be achieved in vivo by delivering a split-intein Cas9 system, sgRNAs, and the donor cassette via adeno-associated virus (hereafter referred to as ACTB-Scarlet REPLACEv1). Although incorporating all components into two AAV vectors enabled REPLACE efficiencies of up to ∼11%, further optimization is needed to reach higher editing levels. Recently, Banskota et. al. introduced engineered virus-like particles (eVLPs) as a novel delivery modality for therapeutic nucleases like Cas9 or base editors ^18^.

These eVLPs are assemblies of viral proteins from murine leukemia virus (MLV) that can encapsulate and release fully assembled Cas9-gRNA or base editor-gRNA ribonucleoproteins (RNPs) directly into target cells, thereby eliminating the need for continuous transgene expression. Because eVLPs lack viral genetic material, they also avoid the risks associated with viral genome integration.

Inspired by this approach, we hypothesized that pairing eVLP-based delivery of Cas9 and gRNAs with AAV-mediated donor delivery could further enhance REPLACE efficiency in vivo. Unlike multiple AAV vectors that may compete for the same receptor, eVLPs and AAVs rely on different entry pathways, thereby minimizing interference and potentially improving co-transduction. Additionally, eVLPs deliver Cas9 transiently, which may help reduce off-target editing.

We then adapted our ACTB-Scarlet REPLACEv1 strategy by replacing the split-intein Cas9 system with eVLPs encapsulating SpCas9 and sgActb-1+2 (hereafter referred to as VLP-Cas9-sgActb-1+2). The ACTB-donor construct was delivered via an AAV vector (Fig. 3a), and this new approach is termed ACTB-Scarlet REPLACEv2. To evaluate its efficacy in vitro, we transduced primary hepatocytes isolated from wild-type mice with both REPLACE components (AAV-ACTB-donor: MOI 500,000 and VLP-Cas9-sgActb-1+2: MOI 250,000) one day after isolation (Fig. 3b). Four days post-transduction, quantification of fluorescence imaging showed that ∼35% (%DAPI) of the total hepatocyte population treated with both AAV-ACTB-donor and VLP-Cas9-sgActb-1+2 displayed Scarlet fluorescence, whereas none of the control samples (untreated, donor-only, or VLP-RNP-only) showed a Scarlet signal (Fig. 3b-c).

**Figure 3.**
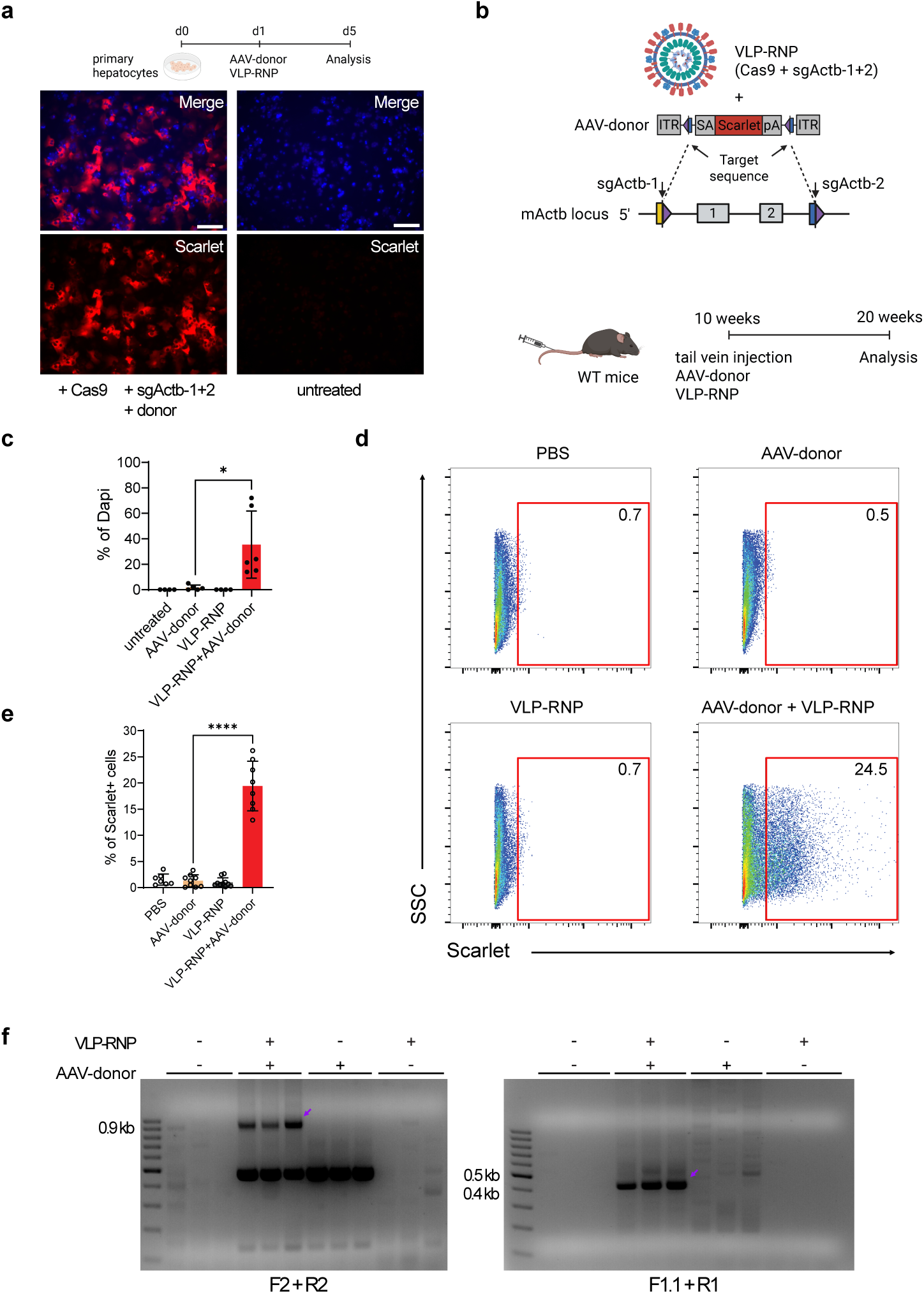
eVLP delivery of Cas9-RNP enhances in vivo REPLACE editing. **(a)** In vitro validation of the ACTB-Scarlet REPLACEv2 strategy. Representative fluorescence images of primary hepatocytes isolated from wild-type mice and transduced in vitro with AAV-ACTB-donor and VLP-Cas9-sgActb-1+2 one day after isolation. Scale bar, 200 µm. **(b)** Schematic representation of the ACTB-Scarlet REPLACEv2 strategy and in vivo experimental workflow. In this configuration, SpCas9 and sgRNAs (sgActb-1 and sgActb-2) are delivered as engineered virus-like particles (VLP-Cas9-sgActb-1+2), whereas the ACTB donor cassette is delivered via AAV D/J. Ten-week-old wild-type male mice were co-injected with both components, and primary hepatocytes were isolated 10 weeks later for analysis. **(c)** Quantification of in vitro editing efficiency shown as the percentage of Scarlet^+^ cells among all hepatocytes (DAPI^+^). Data are mean ± SD from 3 biological replicates (n = 3 independent experiments). **(d-e)** Flow-cytometry analysis of Scarlet-expressing hepatocytes isolated from mice treated with REPLACEv2 strategy compared with control groups (PBS injected, AAV-donor, VLP-RNP). The gates indicate the percentages of ScarletI^+^ cells. Editing efficiency was measured as the percentage of Scarlet^+^ hepatocytes among total hepatocytes. Data are shown as mean ± SD (n = 7-10 mice per group). **(f)** PCR genotyping of FACS-sorted Scarlet^+^ hepatocytes from mice treated with AAV-ACTB-donor and VLP-Cas9-sgActb-1+2. Bulk hepatocytes from control animal groups served as negative controls. Statistical significance was evaluated using an unpaired two-tailed t test with Welch’s correction; * p ≤ 0.05, **** p ≤ 0.0001.

Next, we applied the same strategy in vivo to 10-week-old WT C57BL/6 (B6) mice, which were co-injected with both constructs (AAV: 2 x 10^10^ vg/g mouse, VLP: 5 x 10^9^ vp/g mouse). Ten weeks after injection, primary hepatocytes were isolated from mice and Scarlet expression was quantified by flow cytometry (Fig. 3a, d). Control animals (PBS-injected, AAV-ACTB-donor-only, or VLP-RNP-only) displayed only background levels of Scarlet fluorescence, whereas ∼20% of the total hepatocyte population from mice receiving both AAV-ACTB-donor and VLP-Cas9-sgActb-1+2 were Scarlet-positive (Fig. 3e). This outcome demonstrates that ACTB-Scarlet REPLACEv2 can achieve robust editing in vivo, and can be enhanced by substituting AAVs with VLPs.

To confirm that Scarlet expression reflected correct exon replacement rather than donor integration (HITI) at the sgActb-1 site, we performed PCR-based genotyping as described in Sections 1 and 2. Scarlet-positive hepatocytes were isolated by FACS-sorting from mice treated with AAV-ACTB-donor + VLP-Cas9-sgActb-1+2, and junction PCR was performed on genomic DNA of these cells. We used bulk hepatocytes isolated from control animals (PBS-injected, AAV-ACTB-donor-only, or VLP-RNP-only) as negative controls. Scarlet⁺ hepatocytes displayed the expected ∼930 bp amplicon at the 3′ junction, which is consistent with correct exon replacement at the *Actb* locus (Fig. 3f). In contrast, we did not detect the larger ∼2000 bp product expected for HITI-like integration at the same site. Control samples showed no replacement-specific bands. Together, these genotyping results indicate that the Scarlet signal observed in REPLACEv2-treated animals predominantly arises from correct REPLACE-mediated replacement rather than alternative integration events.

### 4. Large sequence replacement by REPLACE

Although our earlier experiments demonstrate the feasibility of REPLACE for exon-level editing, many research and therapeutic applications require insertion or replacement of larger DNA sequences. A prominent example is the generation of humanized mouse models, in which a mouse gene is replaced entirely or partly with its human homolog, to better model human disease and facilitate the testing of therapeutics in vivo.

To investigate whether REPLACE supports large-scale genomic substitution, we designed hAce2-REPLACE targeting the endogenous mouse *Ace2* locus for replacement with the human *ACE2* coding region. Unlike the ACTB-Scarlet design, this strategy replaces a multi-kilobase interval (∼27kb) spanning mouse *Ace2* exons 1-9 (Fig. 4a).

**Figure 4.**
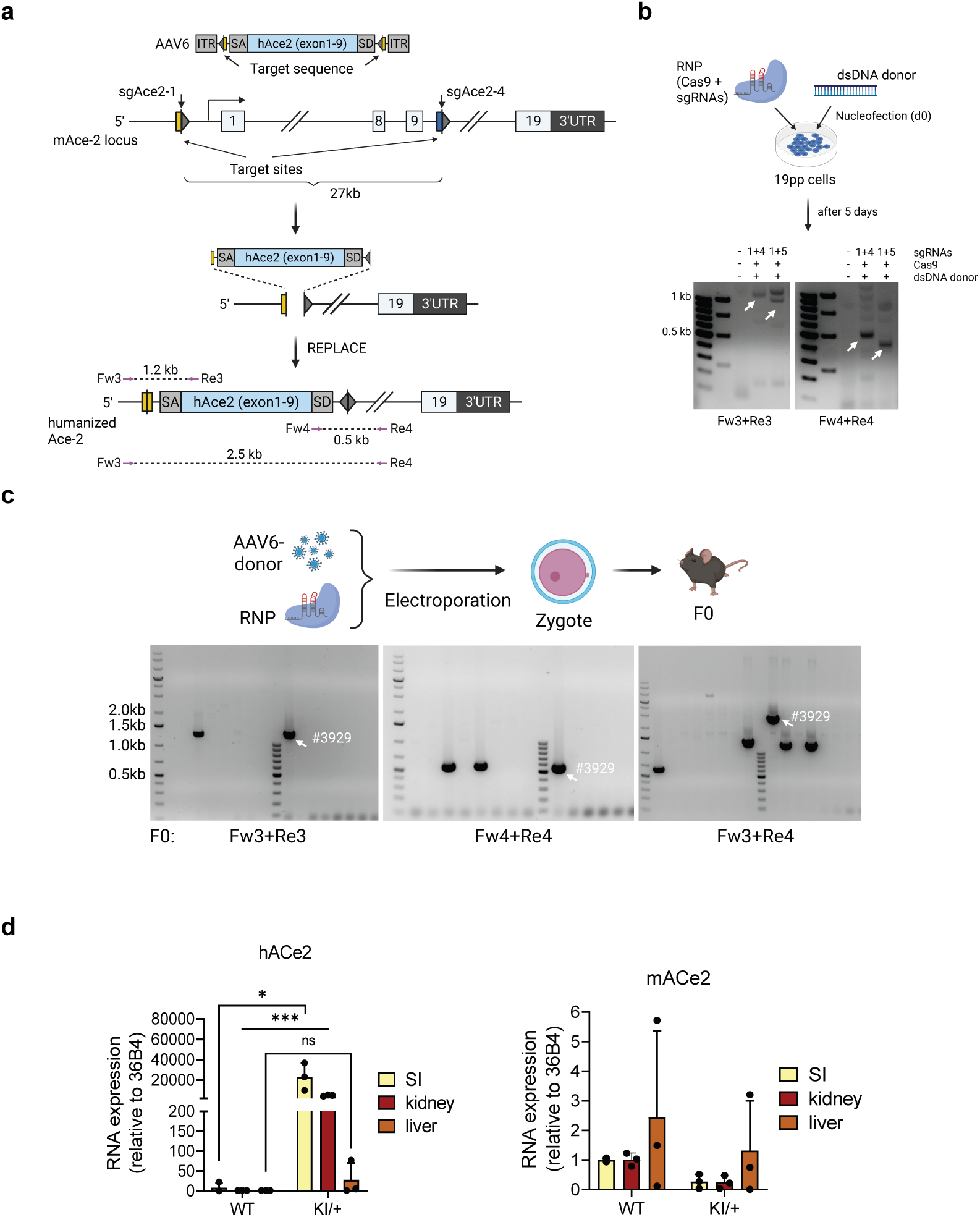
REPLACE supports multi-kilobase humanization of the mouse Ace2 locus. **(a)** Schematic representation of the hAce2-REPLACE strategy. A ∼27-kb genomic interval spanning mouse Ace2 exons 1-9 was targeted for replacement with the human ACE2 coding region (exon1-9). sgRNAs (sgAce2-1 & sgAce2-4) flanking the mouse locus are indicated by black arrows. Genotyping primers are shown as purple arrows and labeled Fw3, Fw4, Re3, and Re4. **(b)** Validation of sgRNA/donor combinations for hAce2-REPLACE in 19PP cells. Cas9/gRNA RNPs and double-stranded donor DNA were delivered by nucleofection, and replacement-specific junction PCR was performed to identify productive combinations. White arrows indicate PCR products consistent with correct replacement. **(c)** Experimental scheme for generation of humanized Ace2 mice. Cas9-gAce2-1+4 RNPs together with AAV6-packaged Ace2-REPLACE donor 1 were delivered by electroporation into C57BL/6 zygotes. Founder animals were screened by junction PCR using the indicated primers. White arrows indicate PCR products consistent with correct replacement. **(d)** RT-qPCR analysis of hACE2 and mAce2 transcript expression in heterozygous hAce2 knock-in (KI/+) and wild-type (WT) littermates. Tissues like small intestine (SI), kidney and liver were analyzed. Data are shown as mean ± SD (n = 3 mice per genotype). Statistical significance was evaluated by unpaired two-tailed t test with * p ≤ 0.05, *** p ≤ 0.001.

We designed 7 sgRNAs flanking the mouse *Ace2* interval, two “upstream” guides (sgAce2-1, -2 upstream exon 1) and five “downstream” guides (sgAce2-4, -5, -7, -11, -12 in intron 9-10) (Supplementary Fig. 4a). All sgRNA upstream/downstream combinations were evaluated in N2A cells using the dual-cut excision assay described in Results Section 1. Cells were co-transfected with plasmids expressing SpCas9 and each sgRNA pair, and the locus spanning both cut sites was PCR-amplified. As expected, no amplicon corresponding to the unedited allele was obtained, because the size of the fragment (> 27kb) is not amplifiable under standard conditions. Productive dual cutting generated one fragment at size ∼ 1-1.5 kb consistent with the excision of exon 1-9 (Supplementary Fig. 4a). All sgRNA pairs showed detectable activity. We next constructed three ACE2-REPLACE donors comprising of the human Ace2 coding sequence corresponding to exon 1-9 flanked by inverted gRNA binding sites of either sgAce2-1, -2 or -11. REPLACE test experiments were performed in 19PP cells nucleofected with Cas9/gRNA RNPs together with double-stranded donor DNA (Supplementary Fig. 4b). Edited cells were genotyped using junction PCR with one primer binding inside the donor and one binding in the genomic locus. Most combinations did not yield replacement-specific products. However, sgAce2-1+4 and sgAce1-1+5 paired with ACE2-REPLACE donor 1 produced the expected replacement amplicons (Supplementary Fig. 4b, Fig. 4b). Notably, the sgAce2-1/4 pair has the feature to reconstitute the sgAce2-1 target sites in several undesired outcomes, enabling iterative re-cleavage and biasing the repair towards correct replacement. Based on this characteristic and the 19PP results, we selected sgAce2-1+4 with ACE2-REPLACE donor 1 for generating the humanized ACE2 mouse.

For production of the humanized allele, the ACE2-REPLACE donor 1 was packaged in AAV6. Cas9-sgRNA RNPs and the AAV6 donor were delivered by electroporation into zygotes isolated from C57BL/6 mice (Fig. 4c). Across several rounds of electroporation, 64 pups were obtained and genotyped by junction PCR using primer pairs as indicated in Figure 4a, c. Among these offspring, 38 animals still carried the wild-type allele (no edit), whereas 19 carried a deleted allele resulted from dual excision without integration of the donor. Five pups carried the donor because of single site integration (HITI) and two animals showed correct full replacement of the human donor across exon 1-9 (Fig. 4c). Sanger sequencing of the correct-replacement founders (mouse #3929) confirmed precise humanization of the *Ace2* allele (Supplementary Fig. 4c).

Positive founder animals were subsequently mated to C57BL/6 WT mice to establish the humanized ACE2 mouse line (here referred to as hACE2 mice). Approximately half of the offspring from the founders carried the humanized *ACE2* allele, therefore confirming germline transmission.

To determine whether the humanized *ACE2* allele was transcriptionally active, we analyzed human ACE2 (hACE2) and endogenous mouse Ace2 (mAce2) mRNA expression by RT-qPCR in heterozygous knock-in (KI/+) and WT littermates (Fig. 4d). We examined tissues with reported high endogenous Ace2 expression, including small intestine and kidney, as well as liver, which expresses Ace2 at lower levels ^19^. As expected, hACE2 transcripts were detected in KI/+ animals but were absent from WT controls, confirming expression of the humanized allele (Fig. 4d). In parallel, mAce2 expression was readily detected in WT animals, whereas its level was reduced in KI/+ mice, consistent with replacement of one endogenous *Ace2* allele. The tissue distribution of hACE2 expression mirrored that of endogenous Ace2, with highest expression in small intestine and kidney and lower expression in liver. These findings demonstrate that the humanized *ACE2* allele generated by REPLACE is transcriptionally active in physiologically relevant tissues.

Together, our results demonstrate that REPLACE can support multi-kilobase humanization of a mouse locus in a single step, using Cas9-RNP electroporation and an AAV6 donor in zygotes. This example shows REPLACE ability extends beyond exon-scale edits to gene-scale substitutions suitable for building humanized translational mouse models.

### 5. REPLACE corrects a frameshift mutation in patient derived cardiomyocytes

We next tested the ability of REPLACE to correct a pathogenic mutation in post-mitotic human cardiomyocytes derived from patients with dilated cardiomyopathy. To this purpose, we targeted a frameshift mutation (K117fs) in the human Lamin gene (*LMNA*) caused by a single-base G insertion at the coding nucleotides 348–349 (348–349insG) in exon 1 ^20^. This heterozygous mutation leads to Lamin A/C haploinsufficiency, thereby activating the PDGF signaling pathway and leading to arrhythmia ^21^ . We used patient-derived induced pluripotent stem cells (iPSCs) engineered to carry the homozygous K117fs mutation and differentiated them into cardiomyocytes, here referred to as hi-CMs. Hi-CMs obtained by in vitro differentiation expressed cardiac markers like Troponin T, the sarcomeric marker alpha-actinin and were negative for the proliferation marker Ki67, indicating their post-mitotic fate and functional character (Supplementary Fig. 5).

To enable the replacement of the mutated exon 1 by REPLACE, we designed six sgRNAs targeting intronic sequences flanking *LMNA* exon 1, three “upstream” guides (sgLMNA-12, -13, -14 in intron 1) and three “downstream” guides (sgLMNA-15, -16, - 17 in intron 2) (Supplementary Fig. 6a).

We evaluated all nine upstream/downstream combinations in HEK293T cells using the simple excision assay as described in Results Section 1. Cells were co-transfected with plasmids expressing SpCas9 and each sgRNA pair. The genomic interval spanning the two cut sites was PCR-amplified. In controls, a single amplicon (WT band: 1.7 kb) corresponding to the unedited locus was observed. Productive dual cutting generated a shorter fragment (600-800 bp) consistent with exon 1 excision. All pairs yielded detectable excision, which was verified by Sanger sequencing of the ligation sites (Supplementary Fig. 6b-c). Importantly, sgLMNA-pair 14+16 is also predicted to reconstitute the sgLMNA-14 target site in several undesired outcomes (e.g., simple religation or inverted donor insertion without indels), allowing iterative re-cleavage of undesired ligations and thereby enrichment of the desired replacement. Based on the strength of these features, sgLMNA-14 and sgLMNA-16 were used for editing in hi-CMs.

The LMNA-REPLACE repair construct consists of a wild-type copy of exon 1 flanked by inverted sgLMNA-16 binding sites and has a length of 800 bp (Fig. 5a).

**Figure 5.**
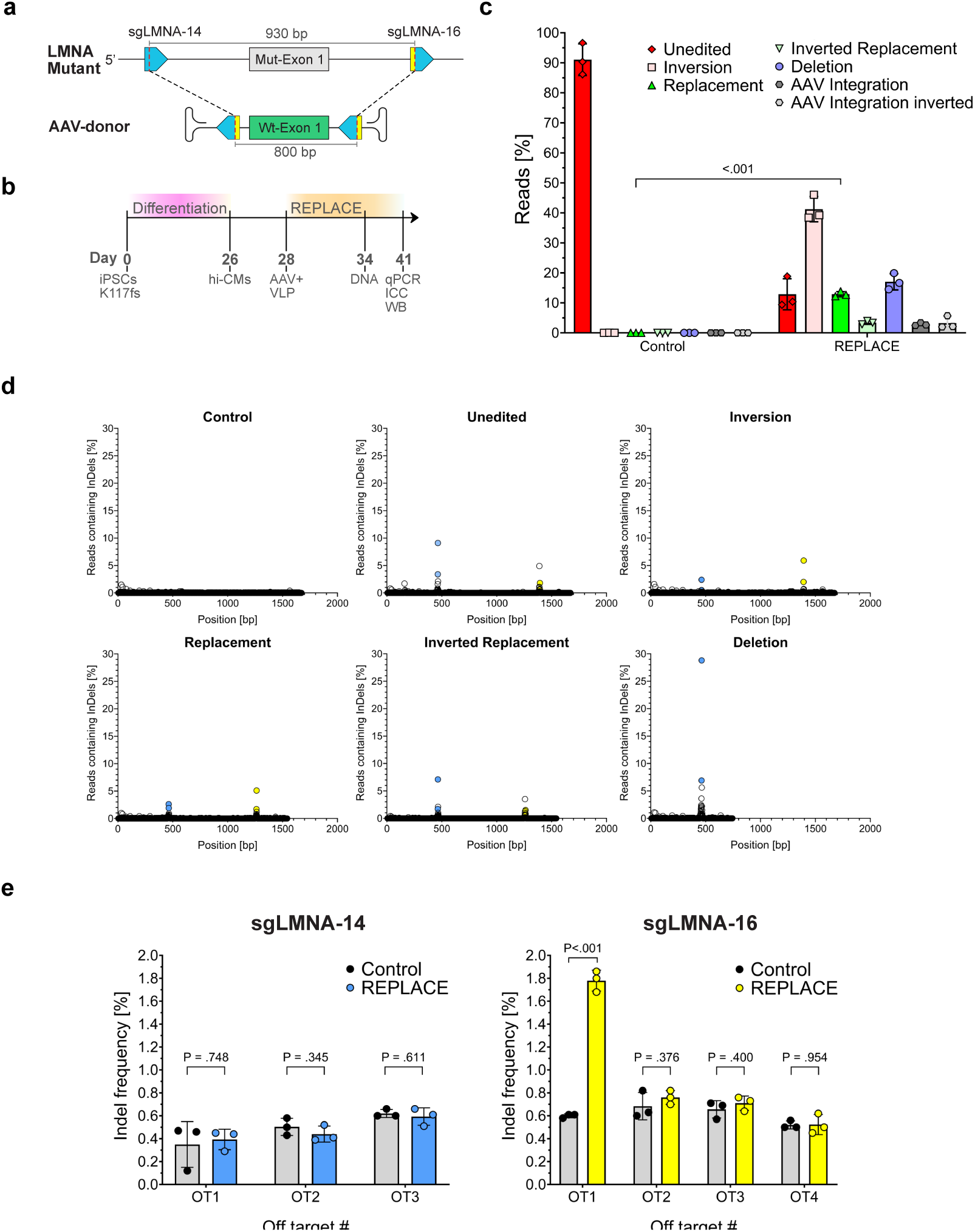
REPLACE corrects the human LMNA K117fs mutation in post-mitotic human cardiomyocytes. **(a)** REPLACE strategy targeting exon 1 in the human LMNA gene. **(b)** Experimental timeline of REPLACE-editing in hi-CMs. **(c)** Quantification of genomic structural variants in untreated control and REPLACE-treated hi-CMs by Long-read PacBio sequencing. **(d)** Indel frequencies [%] across the whole Amplicon length of the respective structural variant obtained by Long-read PacBio sequencing. Indel rates 1 bp around the gRNA cutting sites are labelled in blue for sgLMNA-14 and in yellow for sgLMNA-16. **(e)** Indel frequency [%] at in silico predicted off-target (OT) sites of sgLMNA-14 (left) and sgLMNA-16 (right) quantified by Illumina Amplicon sequencing. Data is expressed as mean ± SD from three biological replicates (n=3 different AAV- and VLP-batches). Unpaired two-tailed student’s t-test was performed.

On day 28th of differentiation, hi-CMs were incubated with SpCas9 and selected gRNAs (sgLMNA-14+16), which were co-delivered in eVLPs (1.03 x 10^4^ vp/cell). The LMNA-REPLACE donor construct was delivered in an AAV-D/J vector at 3 x 10^6^ vg/cell at the same time. After six days, we quantified editing outcomes, indel and off-target frequencies by deep sequencing (Fig. 5b).

To quantify correct exon replacement and alternative outcomes (e.g. deletions and inversions), we performed long-read sequencing (PACBIO) across the entire edited interval using primers binding upstream of the 5’ and downstream of the 3’ cut site.

Among all reads, ∼13% corresponded to the expected exon replacement event with WT exon 1 in the correct orientation. We observed comparable frequencies of unedited (13%) and deletion (17%) events. Notably, ∼ 41% of the reads contained an inversion of the endogenous mutant exon 1, arising from dual cutting and reinsertion of the excised fragment in the inverted orientation at the target locus. By contrast, inverted replacement (REPLACE with inverted wt-donor, 4%) or AAV backbone integration (5.5%) were rare (Fig. 5c).

To better resolve the distribution of the different editing variants after REPLACE treatment, we quantified indel frequencies of REPLACE treated samples within ±1 bp of each sgLMNA cut site. Correct exon replacements carried indels at the sgLMNA-14 and -16 cut sites in 4.5% and 6.8% of reads, respectively, whereas alleles containing the endogenous exon fragment in the correct orientation, here referred to as unedited alleles, harbored 12.5 and 2.5% indels at the same sites. Deletion alleles showed the highest burden with 35.7% at the re-ligation site. Endogenous inversions exhibited indels at 2.9% (sgLMNA-14) and 7.9% (sgLMNA-16), while inverted donor replacement carried 8.8% and 2.1% indels at sgLMNA-14 and -16, respectively (Fig. 5d).

To assess off-target activity, we used CRISPOR ^22^ to select candidate sites *in silico* and analyzed seven sites by targeted amplicon sequencing in untreated and REPLACE-treated hi-CMs. Six of seven OT sites showed no significant increase in indel frequency relative to baseline. Only OT#1 for sgLMNA-16 exhibited a threefold increase over control to 1.8%, which is still very low in absolute terms (Fig. 5e).

The molecular effect of REPLACE-mediated correction of the K117fs mutation in homozygous hi-CMs was analyzed 13 days after transduction (Fig. 5b). Western blotting of the Lamin A/C proteins encoded by the LMNA gene showed no Lamin A/C protein expression in untreated hi-CMs carrying the homozygous K117fs-mutation, whereas REPLACE-treated cells re-expressed the full-length Lamin A/C matching the expected size and of the isogenic wild-type hi-CMs (Fig. 6a). Notably, edited samples reached 18.8% of isogenic wild-type protein levels, reaching a 28.5-fold increase over untreated controls (P=.026, Fig. 6b). Furthermore, the expression levels and cellular localization of Lamin A/C were analyzed by immunostaining. Confocal imaging of stained hi-CMs revealed correct localization of Lamin A/C in the nuclear lamina upon REPLACE-treatment, while untreated mutant hi-CMs showed no Lamin A/C expression (Fig. 6c). Quantification of Lamin A/C fluorescence intensity for each single-nucleus revealed a significant increase in REPLACE-treated hi-CMs compared with untreated mutant controls (P<.001, Fig. 6d). The mean Lamin A/C signal intensity per nucleus was 2.3-fold higher after REPLACE treatment and reached approximately 52% of the signal detected in isogenic wild-type controls (Fig. 6d). Overall, these results show that REPLACE-editing of the K117fs-mutation in hi-CMs restores Lamin A/C protein expression and nuclear lamina localization in post-mitotic human cardiomyocytes.

**Figure 6.**
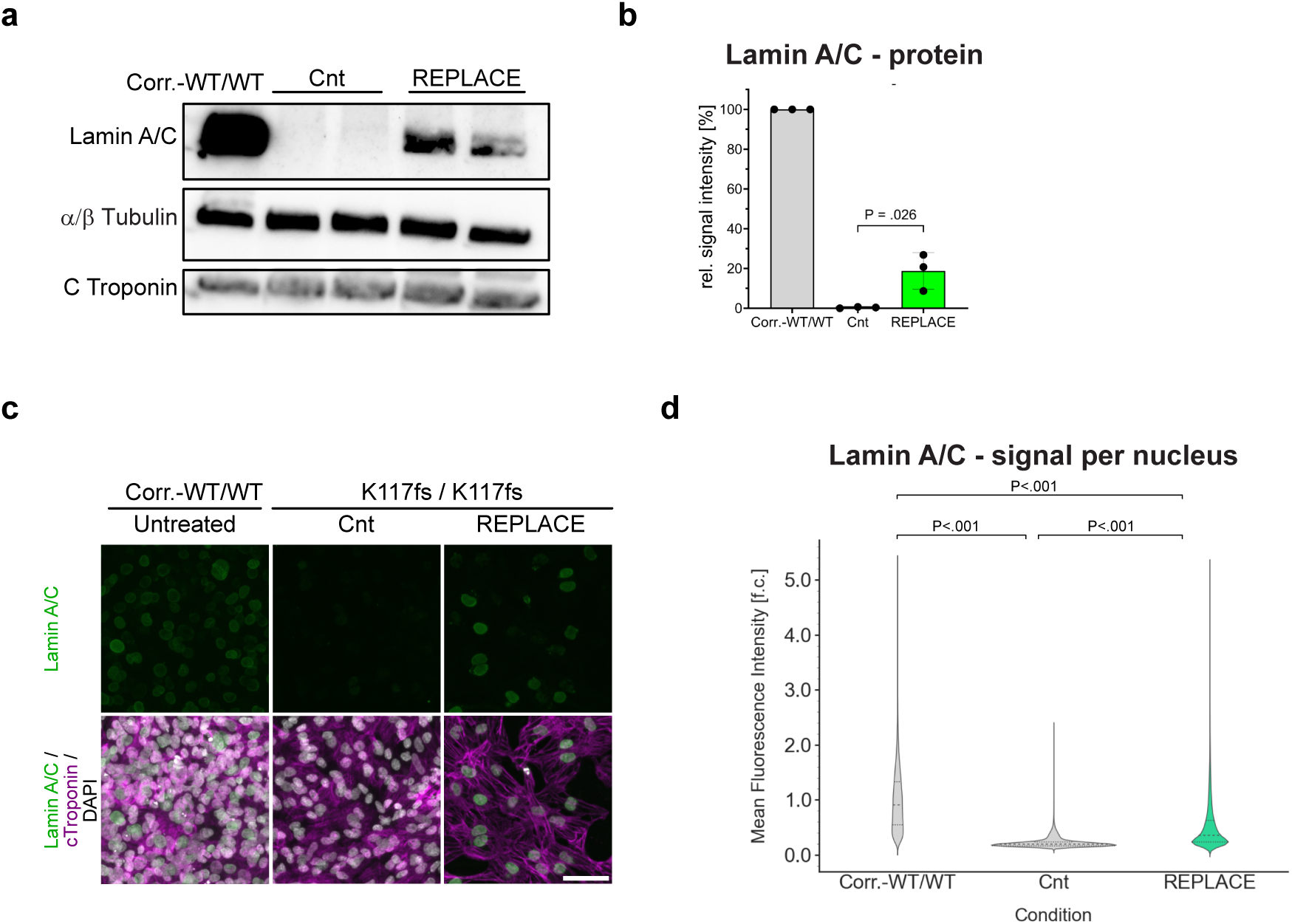
REPLACE restores LMNA protein expression in edited K117fs mutants. **(a)** Representative Western Blot image of Lamin A/C protein in hi-CMs. α/β-Tubulin and c-Troponin were used as loading controls. **(b)** Quantification Lamin A/C band intensity shown in a. Values were expressed as fold change [f.c.] relative to isogenic wildtype control, which was set to 100%. Data are shown as mean ± SD from three biological replicates (n=3 independent differentiations). Statistical significance was determined using an unpaired two-tailed Student’s t test. **(c)** Representative confocal microscopy images of fluorescence-stained isogenic wildtype, untreated mutant, and REPLACE-treated mutant hi-CMs. DAPI (white), Lamin A/C (green) and cTroponin (magenta). Scale bar, 50 μm. **(d)** Quantification of Lamin A/C fluorescence intensity at the single nucleus level based on confocal images aquired with a 20× objective. At least 42,000 nuclei were analyzed per condition from three independent differentiations. Values are expressed as fold change [f.c.] relative to the isogenic wild-type control, which was set to 1. Statistical significance was determined using an unpaired two-tailed t-test with Welch’s correction. Violin plots show the 25th and 75th percentiles as dotted lines and the median as a dashed line.

## Discussion

CRISPR-Cas9-based gene correction holds great potential for gene therapy ^23^. Despite intensive development and improvement, precise sequence replacement in non-dividing cells remains a central hurdle for therapeutic applications. Here, we present REPLACE as a general gene replacement approach that is applicable in primary cells, in vivo tissues, and human post-mitotic cardiomyocytes. Across diverse settings, REPLACE has led to precise exchange of defined genomic segments.

In Cas9-expressing primary hepatocytes, REPLACE achieved high replacement among transduced cells (∼74%), proving that NHEJ-mediated exon replacement can be efficient in primary cells. In the adult liver, split-intein AAV delivery of Cas9 with an AAV donor produced ∼11% correctly edited hepatocytes, providing a baseline for in vivo replacement. To improve in vivo replacement efficiency, we decoupled nuclease and donor delivery by using different delivery modalities. This strategy can alleviate co-transduction limits by leveraging non-competing entry pathways. Building on recent eVLP technology, we combined eVLP-delivered Cas9-gRNA RNPs with AAV-delivered donor. This hybrid delivery approach increased in vivo editing significantly (∼20% Scarlet^+^ cells among total hepatocytes).

Another important aspect of this work is the ability of REPLACE to install multi-kilobase large substitutions. Using RNP electroporation with an AAV6 donor, we replaced a ∼27-kb large segment of the mouse *Ace2* locus spanning exons 1-9 with the human *ACE2* ortholog in zygotes. Genomes of founder mice spanned the expected spectrum of editing outcomes (WT, deletion-only, HITI-single site integration), including the precise full replacement. While the full-replacement rate was modest (∼3.1%), the experiment underscores the applicability of REPLACE for gene-scale humanization in a single step. Thus, REPLACE constitutes an alternative HDR-independent technology to generate challenging translational knock-in models.

Previously, large-segment humanization, e.g., insertion of the full human *ACE2* gene of ∼116–180 kb, has been achieved with the genome-writing method mSwap-In in mouse embryonic stem cells (ES cells) ^24^. While mSwap-In is a powerful technology to achieve full-length genomic humanization (including all non-coding/regulatory regions), it requires donor assembly in yeast and multistep, iterative donor integrations in ES cells prior to chimera generation. The complexity of the editing process renders the workflow comparatively labor-intensive and technically challenging. By contrast, REPLACE works as a one-step, single-embryo procedure, enabling relatively rapid model generation. Owing to its shorter workflow and the reduced experimental complexity, REPLACE may represent a more practical approach in case of humanization of the coding sequence (that is, lacking the non-coding regions of the human gene) is required.

Finally, we applied REPLACE to correct a clinically relevant frameshift-mutation (K117fs) in the human *LMNA* gene in post-mitotic hi-CMs to evaluate the feasibility of REPLACE as a therapeutic strategy. REPLACE-editing in hi-CMs yielded robust 13% correct exon replacement among long-read sequences, with low off-target activity (< 2%). Using eVLPs to deliver Cas9-sgRNA RNPs and AAV for the donor, REPLACE restored Lamin A/C protein expression in homozygous mutant hi-CMs to 18.8% (Western Blot) or 52% (immunostaining) of isogenic wild-type levels.

The two sgRNAs flanking *LMNA* exon 1 were selected based on their identical sequence at the 3’ end in order to minimize unwanted editing outcomes. Long-read sequencing across entire exon 1 and both gRNA-cut sites confirmed donor integration in the desired orientation at an approximately 3:1 ratio to its inverted integration, indicating that rational gRNA/donor design can direct donor integration without the use of homology arms.

Nevertheless, comprehensive long-read profiling of the *LMNA* locus revealed that REPLACE results in a number of alternative editing outcomes in addition to the correct exon replacement. The most frequent alternative outcome was the inversion of the endogenous exon 1 (∼41%), which likely arises from rapid re-ligation of the excised endogenous fragment before donor capture. Because the inverted re-insertion of the mutant exon does not recreate the sgLMNA-recognition sites, these edited alleles are impervious to iterative re-cleaving and accumulate in the cell population. Moreover, we observed increased rates of alleles harboring a deletion between the two cut sites (17%). High indel-rates at the sgLMNA-14 cut site (35.7%) indicate that disruption of the gRNA binding site after initial cleavage prevented further targeting, leading to an accumulation of the deletion product.

The fraction of undesired editing outcomes, such as inversions and large deletions, should be minimized in further studies by refining sgLMNA selection. Given that the indel-rates of sgRNAs depend on the cut site, its flanking regions and the sgRNA sequence itself, the mutational profile of sgRNAs is predictable and reproducible ^25,26^. Therefore, the flexibility to position the REPLACE-sgRNAs almost anywhere within the intronic region can be harnessed to select sgRNAs with inherently lower indel profiles to preserve the gRNA targeting sites for re-cleavage. Another factor influencing the purity of correct exon replacement is the blunt-end cutting nature of SpCas9. To prevent the uncontrolled re-ligation of genomic and plasmid blunt DNA ends, staggered-cut inducing nucleases, such as Cas12a or Cas12e ^27,28^, should be evaluated to improve the REPLACE technology. The 5’-overhangs create a unique sequence, allowing directed re-ligation of the genomic DNA ends with the donor sequence, thereby preventing exon inversions and deletions.

Targeted amplicon sequencing at CRISPOR-nominated off-target sites revealed no significant increases at six out of seven sites, while one site (OT 1 of sgLMNA-16) exhibited a slightly elevated indel-frequency (1.8%) in REPLACE-treated hi-CMs.

This OT site is located within an intron in the Coiled-coil domain-containing protein 38 (CCDC38), a gene implicated in spermiogenesis. Given the low indel frequency and the intronic location of the OT site, the potential functional risk is estimated to be low. However, comprehensive genome-wide off-target profiling, assessment of chromosomal rearrangements, and evaluation of p53-mediated DNA damage responses will be important before therapeutic application of REPLACE ^29–31^.

In the recent years, next-generation CRISPR tools, such as base and prime editing have been published, which benefit from an HDR-independent, single-nick based working mechanism ^9,10^. Nevertheless, their wide-spread application is limited, as base editing is restricted to introducing single base transversions and the editing efficiency of prime editing remains unpredictable.

In a previous study, our group has established a prime editing strategy in hi-CMs targeting the homozygous K117fs mutation. While the mutation could be corrected in HEK293T cells (37.3%), the same strategy did only yield low editing efficiencies in homozygous hi-CMs (1%) ^11^. Given that PE was ineffective in hi-CMs and base editing is inherently unsuitable for the correction of insertion-type mutations, REPLACE-editing offers the first gene-correction approach of the LMNA^K117fs^-mutation resulting in Lamin A/C re-expression, despite the higher risks associated with dual-cleavage compared to single-nicking methods.

Together, our findings establish REPLACE as a versatile homology-independent genome-editing platform that expands NHEJ-based editing beyond disruption and exon skipping towards precise exon- and gene-scale sequence replacement. By addressing central limitations of current genome-editing technologies, REPLACE extends the therapeutic and translational potential of CRISPR-based editing to genetic contexts in which HDR, base editing, or prime editing are inefficient or not applicable. This work therefore opens new opportunities for somatic gene correction, disease modeling, and the generation of humanized animal models.

## Methods

### Construction of plasmids

Synthetic sgRNA-oligonucleotides (Table 1) were ligated into BbsI-digested pU6-plasmids. Dual-sgRNA-plasmids were constructed using Gibson Assembly. The donor plasmid was cloned by digesting the synthetic donor casette (IDT) and the single-stranded pAAV backbone with restriction enzymes MluI and PmlI. Correct insertion of the donor and integrity of inverted terminal repeats was confirmed by whole plasmid sequencing (Eurofins). Key plasmids used for REPLACE in this study will be deposited with Addgene and made available upon publication.

**Table 1.**
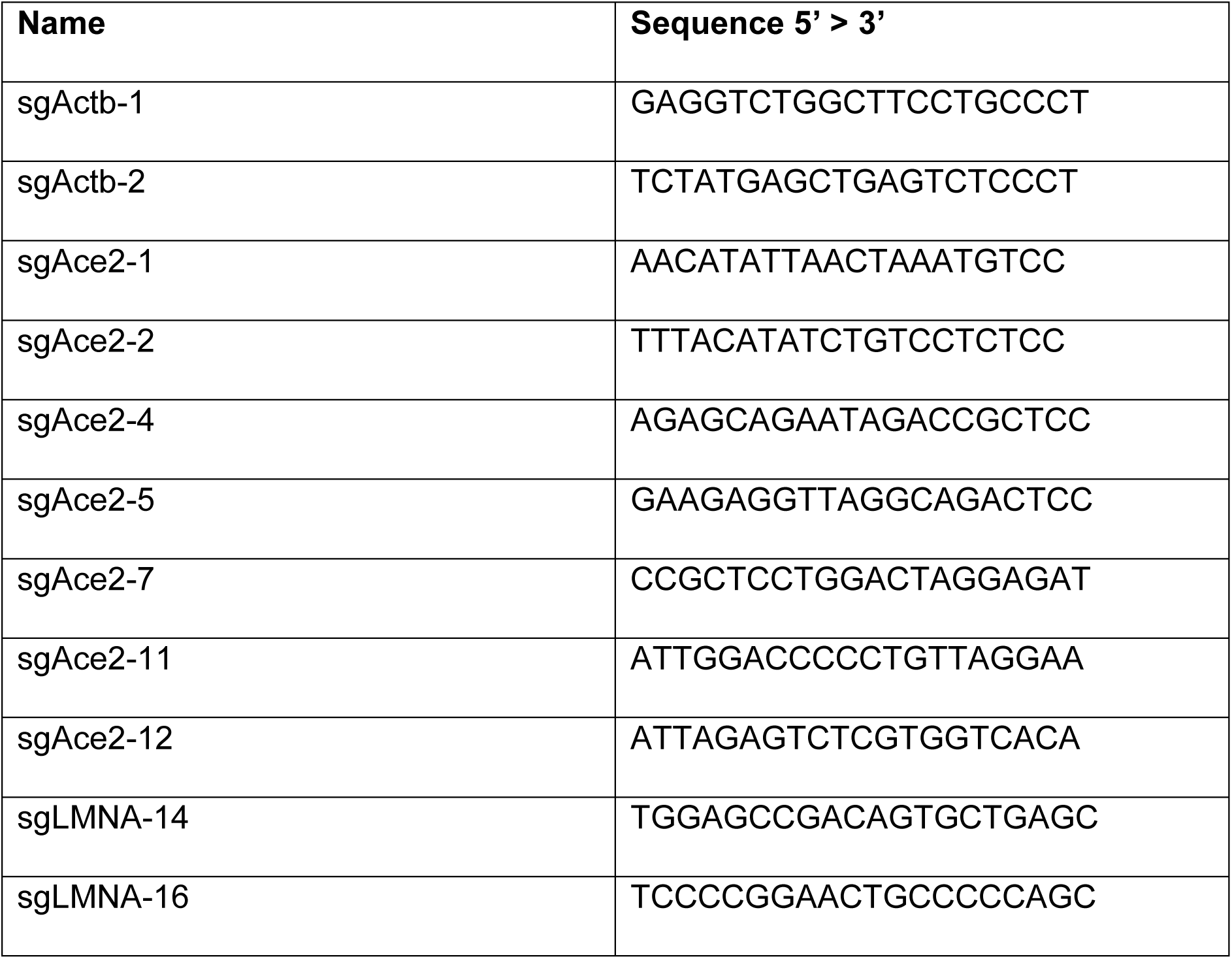
gRNAs.

### Animals

All mice were on a pure C57BL/6N background. Mice were housed in a pathogen-free animal facility at the Max Delbrück Center (Berlin-Buch). Wild-type C57BL/6N, and R26-Cas9 mice were bred in house. R26-Cas9 mice were reported previously ^32^. R26-Cas9 and wild-type mice were injected with either AAV (2 x 10^10^ vg/g mouse for each construct) or together with eVLP (5 x 10^9^ vp/g mouse) particles by tail vein injection. All animal experiments were approved by the Institution Animal Care and Use Committee (Berlin LaGeSo).

### Generation of hAce2 mice

sgRNAs were designed using CRISPOR to target intronic regions flanking the mouse *Ace2* genomic interval spanning exons 1-9. Based on preliminary excision and donor-validation experiments in cultured 19PP cells, sgAce2-1 and sgAce2-4 were selected for further experiments. The donor construct (ACE2-REPLACE donor 1) contained the human *ACE2* coding sequence corresponding to exons 1-9 and was flanked by inverted sgRNA target sites and packaged into AAV6 particles. Zygotes obtained from superovulated C57BL/6N females (Charles River, Sulzbach, Germany) were electroporated as described ^33^ with sgRNA/Cas9 RNPs and AAV6 donor (3.1 x 10^8^ vg/µL). Following electroporation, embryos were cultured to the appropriate stage and transferred into pseudo-pregnant recipient females to obtain mutant founder mice. To establish the hAce2 mouse line, founder males carrying the correctly replaced allele were crossed with wild-type C57BL/6N females.

### Primary hepatocyte isolation

The primary hepatocyte isolation protocol was adapted from the previously described methods by Zhang et al. and Denzler et al. ^34,35^. Male C57BL/6N mice aged 8–12 weeks were euthanized by inhalation of isoflurane. The liver was perfused by cannulation of the caudal vena cava, with the portal vein used as an outlet. First, the liver was perfused with Hank’s Balanced Salt Solution (HBSS; Life Technologies) containing 0.5 mM EGTA, followed by pre-warmed digestion medium consisting of low-glucose DMEM (1 g/L glucose; Life Technologies) supplemented with 30 µg/mL Liberase TM Research Grade (medium thermolysin concentration, Roche). Each perfusion step was carried out for 4 min at a flow rate of 3 mL/min. After perfusion, the liver was surgically removed, and hepatocytes were released into 10 mL digestion medium by gentle shaking. The suspension was supplemented with 15 mL ice-cold low-glucose medium consisting of low-glucose DMEM (1 g/L glucose; Life Technologies), 1% penicillin-streptomycin (Life Technologies), 10% heat-inactivated fetal bovine serum (Sigma), and 1% GlutaMAX (Life Technologies), and subsequently filtered through a 100 µm cell strainer (BD). The cell suspension was washed three times with 25 mL ice-cold low-glucose medium by centrifugation at 200×g for 3 min at 4°C. Hepatocytes were counted and seeded at a density of 300,000 cells per well in surface-treated 6-well plates (Corning Primaria) containing low-glucose medium. Six hours after plating, the medium was replaced with HepatoZYME medium consisting of HepatoZYME-SFM (Life Technologies) supplemented with 1% penicillin-streptomycin and 1% GlutaMAX. Fourteen hours after plating, cells were transduced with AAV and/or VLP particles as indicated. All cells were maintained at 37°C in a humidified incubator with 5% CO₂.

### eVLP production

eVLPs were produced as described previously ^18^. Briefly, 5 x 10^6^ gesicle producer HEK cells were co-transfected with plasmids expressing VSV-G (800 ng, Addgene #8454), MMLV Gag-Pro-Pol (6.75 µg, Addgene #35614), MMLV Gag-SpCas9 (2,25 µg, Addgene #181753), sgRNA-A (8.8 µg) and sgRNA-B (8.8 µg) using polyethyleneimine (PEI, DNA:PEI ratio 1:3, Polysciences). Three days after transfection, the cell supernatant was passed through a 0,45-µm PVDF filter and centrifuged at 2 x 10^5^×g for two hours at 18°C in a Beckman Type 70Ti rotor. The precipitated eVLPs were resuspended in PBS and quantified using a FastScan™ Cas9 ELISA Kit (Cell Signaling).

### rAAV production and transduction

Recombinant AAV (rAAV) was produced as previously described ^36^. Briefly, 5 x 10^6^ HEK-293T cells were co-transfected with 10 μg pAAV-helper, 5 μg pRep/Cap-D/J and 5 µg transgene-encoding plasmid using PEI (Polysciences). rAAV was precipitated from the medium using Polyethylene Glycol (PEG), and extracted from the cells by four rounds of freeze-thawing. Residual DNA plasmids and non-encapsulated viral DNA was removed by benzonase treatment (50 U/mL, VWR International). The solution was purified over an iodixanol gradient (Sigma-Aldrich) of four phases (15%, 25%, 40%, and 60%) and ultracentrifuged at 2 x 10^5^×g for two hours at 18°C in a Beckman Type 70Ti rotor. The rAAV-containing phase was extracted with an 18 g needle and filtered through a 0.22 μm polyethersulfon (PES) syringe filter (Thermo Fisher). Dialysis was performed in a Slide-A-Lyzer Dialysis Cassette (10K MWCO, Thermo Fisher) to exchange the buffer to PBS and to increase the rAAV concentration. Residual DNA was removed via digestion with DNase I (Qiagen). qRT-PCR with a TaqMan probe and primers binding to the Inverted Terminal Repeats (ITRs) was performed to quantify the rAAV titer.

### Cell culture

HEK293T, N2A and Gesicle Producer 293T cells (Takara) were cultured in DMEM+GlutaMax (Thermo Fisher) supplemented with 10% (v/V) FBS and 5% (v/V) Pen/Strep (Thermo Fisher). 19PP cells were cultured in the same base medium additionally supplemented with 5% HEPES (1 M stock), non-essential amino acids (100X), and 52 µM β-mercaptoethanol. All cells were maintained at 37°C in a humidified incubator with 5% CO₂.

### sgRNA tests in HEK293T and N2A cells

Wild-type HEK293T or N2A cells were co-transfected with plasmids expressing dual sgRNAs together with a plasmid encoding SpCas9-T2A-mCherry using PEI. Genomic DNA was isolated 3 days after transfection and used for PCR amplification of the targeted loci.

### RNP and linear DNA electroporation in 19PP cells

For sgRNA complex formation, crRNA and tracrRNA oligonucleotides were mixed at equimolar concentrations (5 µL of 200 µM each), incubated at 95°C for 5 min in a thermal cycler, and then cooled to room temperature. For dual-sgRNA Cas9 RNP assembly, 1.2 µL (120 pmol) of each hybridized crRNA:tracrRNA complex was incubated with 1.7 µL SpCas9 protein (stock concentration 61 µM). 19PP cells were collected in a 15 mL Falcon tube and counted. Cells were washed once with PBS at room temperature, and 0.2 x 10^6^ cells were resuspended in 20 µL Amaxa electroporation buffer. The 4.1 µL total RNP mix was then added together with 400 ng of linear hACE2 donor DNA generated by PCR. Primers used for amplification of the linear donor are listed in Table 2. The final electroporation mixture was carefully transferred into one well of a 16-well Nucleocuvette strip. Electroporation was performed using the Amaxa 4D-Nucleofector (Lonza) with the program specified as CM-120 ^2,37^. Immediately after electroporation, 80 µL of pre-warmed complete medium was added to each well. Cells were then transferred into a 24-well plate containing 1 mL of the same medium. Genomic DNA was isolated 3 days after electroporation.

**Table 2.**
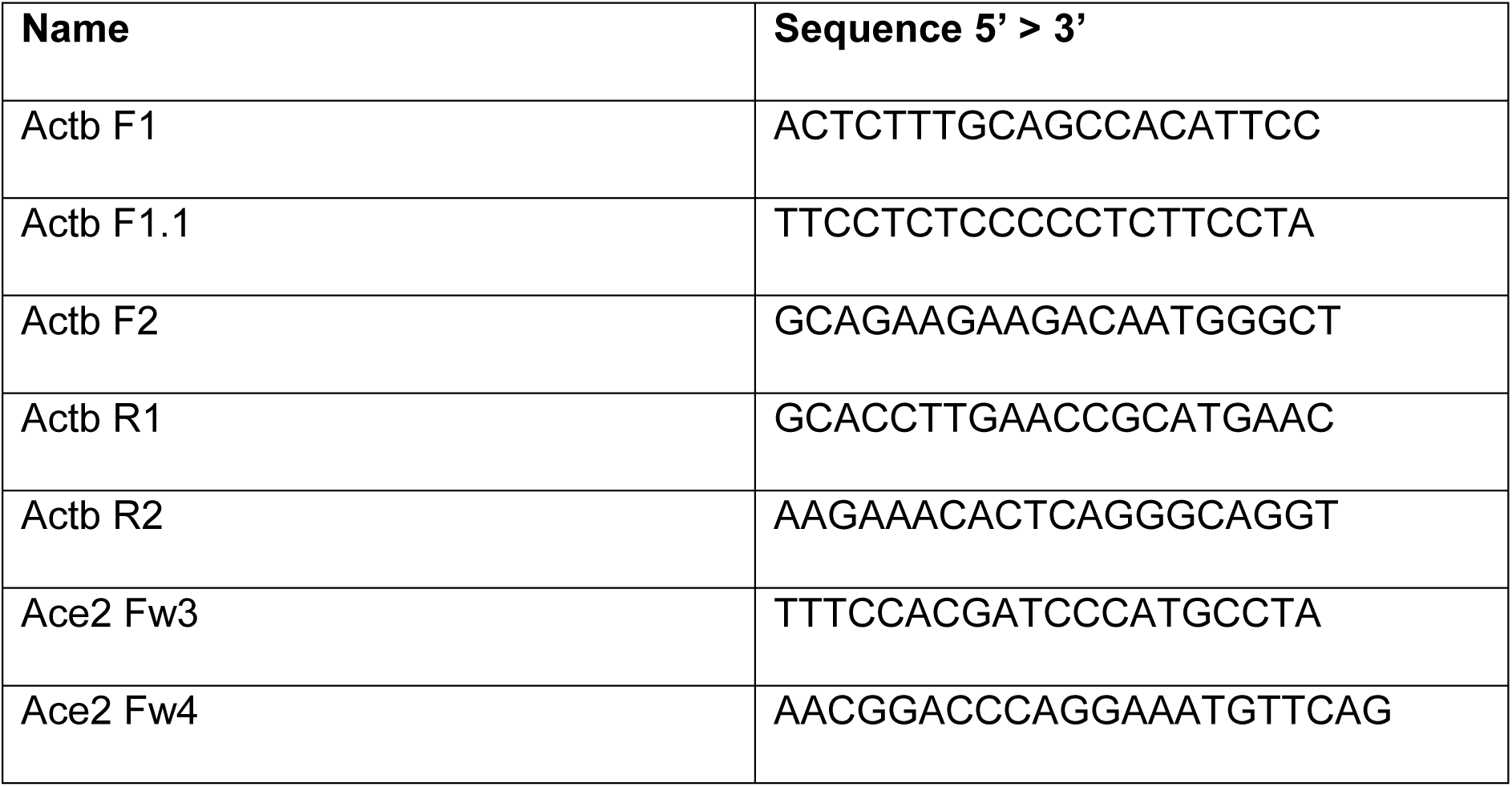
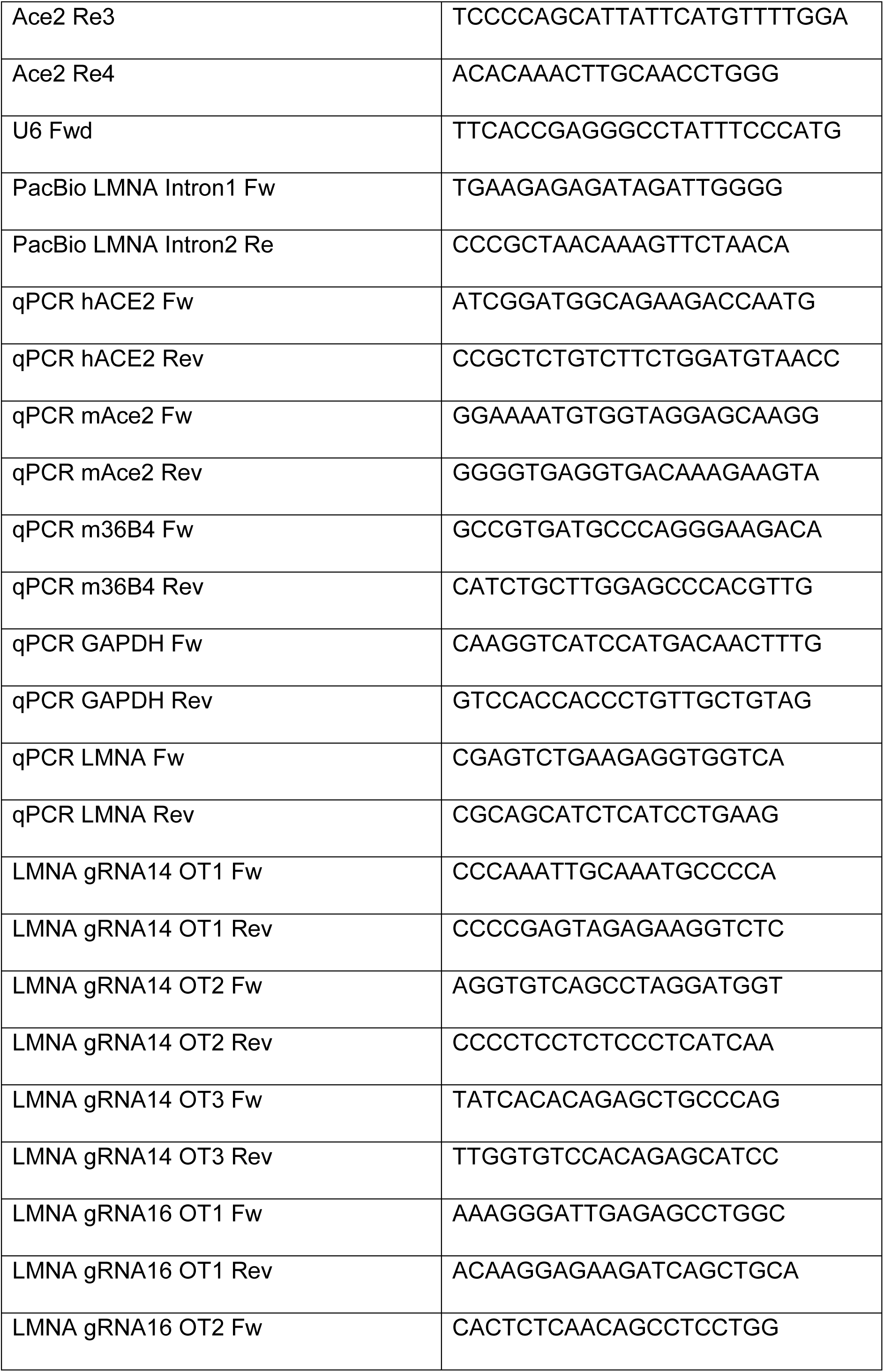
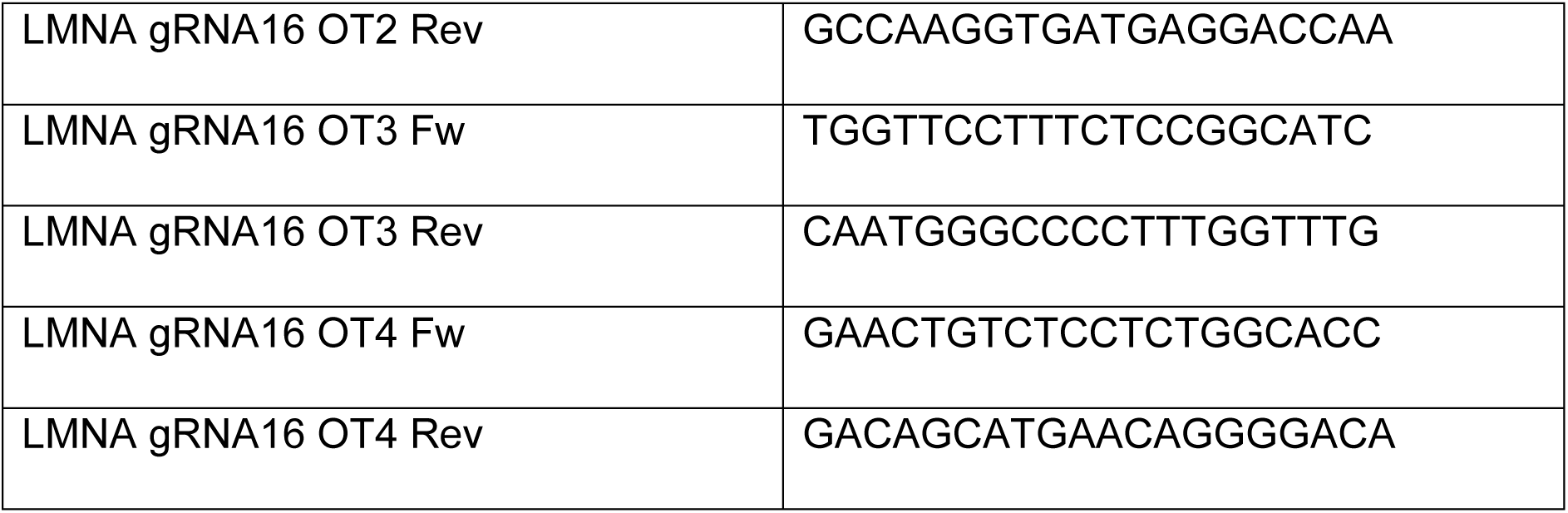
Primers.

### Culture conditions for human iPSCs and cardiac differentiation

Patient-derived iPSCs and their isogenic wild-type control iPSCs were previously generated and published ^21^. iPSCs were cultured on Geltrex (Thermo Fisher) coated plates in Essential 8 Flex (E8, STEMCELL Technologies) medium and passaged with 0.5 mM EDTA/PBS every 2-3 days. At 90% confluence, cardiac differentiation was initiated by changing the medium to RPMI-1640 (Thermo Fisher) supplemented with CHIR-99021 (6-9 μM, Selleckchem) and B27 without Insulin (Thermo Fisher) ^38^. After 24 hours, equal volume of RPMI supplemented with B27-Insulin was added to the cells. At day three, the medium was replaced with RPMI-Insulin and IWR-1-endo (5 μM, Selleckchem). From day seven on, cells were cultured in RPMI with complete B27 supplement (Maintenance medium, Thermo Fisher), which was changed every 2-3 days. Between days 12-15, metabolic selection was performed in RPMI-1640 without glucose (Thermo Fisher), supplemented with sodium DL-lactate (Sigma-Aldrich) and chemically defined medium supplement (CDM3). Hi-CMs were dissociated, frozen and thawed as previously described ^39^. Briefly, hi-CMs were dissociated using 10x TrypLE (Thermo Fisher) and re-plated in maintenance medium supplemented with 10% knock-out serum replacement (KOSR, Thermo Fisher) and RevitaCell supplement (1:100, Thermo Fisher) between day 23 and 25. The medium was changed back to maintenance medium the following day. rAAVs (3 x 10^6^ vg/cell) and eVLPs (1.03 x 10^4^ vp/cell) were added to the hi-CMs at day 28 and incubated for two more days. Cells were harvested for genomic analysis on day 34. RNA and protein analysis was performed on day 41 of differentiation.

### Genome sequencing and data analysis

Genomic DNA (gDNA) was isolated from hi-CMs using the Quick-DNA Miniprep Kit (Zymo Research) and eluted in ddH20. PCR amplification was performed with LongAmp Taq DNA Polymerase (NEB) for On-target and Platinum SuperFi II DNA-Polymerase (Thermo Fisher) for OT analysis. Respective primer sequences are provided in Table 2. Long-read sequencing was conducted on Pacific Biosciences Revio (BIH/MDC Genomics Technology Platform). Next-generation amplicon sequencing was performed at GENEWIZ (Amplicon EZ service) using an Illumina MiSeq platform and 250 bp paired-end reads. Results were analyzed using CRISPResso2.

### RNA isolation and qPCR

Total RNA was isolated from cultured cells or tissue samples using TRIzol (Thermo Fisher) following the manufacturer’s protocol. 900 ng total RNA were reverse transcribed to cDNA using the High-Capacity cDNA Reverse Transcription Kit (Thermo Fisher). qPCR was performed using 2 μL cDNA together with Platinum SYBR Green qPCR SuperMix (Thermo Fisher) following the manufacturer’s instructions in a QuantStudio 7 Pro Real-Time PCR System (Thermo Fisher). Primers binding to reference and target genes are listed in Table 2. C_Q_-values were normalized to the reference gene GAPDH of each sample. Relative gene expression levels were calculated using the formula 2^−ΔΔCQ^.

### Immunocytochemistry of hi-CMs and fluorescence intensity quantification

iPSCs and hi-CMs were grown on IbiTreat-coated 8-well μ-Slides (Ibidi) and fixed with 4% paraformaldehyde (PFA) for 15 minutes. Fixed cells were permeabilized with 0,05% Tween 20 (Sigma-Aldrich) and 0,025% Triton X-100 (Carl Roth) and simultaneously blocked with 1% bovine serum albumin (BSA, Sigma-Aldrich) for one hour at room temperature. Primary antibodies (Table 3) were incubated overnight at 4°C before incubating DAPI and Alexa Fluor-conjugated secondary antibodies at room temperature for 45 minutes. Images were acquired with a confocal TCS SP8 DLS microscope (Leica Microsystems) using a 20x dry objective for fluorescence intensity quantification and a 63x glycerol objective for representative images. Fluorescence intensity was quantified using an ImageJ macro. Briefly, the DAPI channel was used for automated nuclear segmentation with StarDist (model: “Versatile (fluorescent nuclei)”; probability threshold: 0.5; NMS threshold: 0.8; percentile normalization: 1-99.8%). Mean fluorescence intensity of Lamin A/C was measured per nucleus within the resulting ROIs. Values were normalized to the mean of all Corr.-WT/WT nuclei (set to 1.0) and plotted as violin plots using Python (seaborn). Statistical comparisons between conditions were performed using Welch’s t-test.

**Table 3.** Antibodies for immunofluorescence staining and Western Blot.

| Antibody | Species | Dilution | Manufacturer | ID |
| --- | --- | --- | --- | --- |
| Lamin A/C (E-1) | mouse | 1:100 (IF); 1:500 (WB) | Santa Cruz | sc-376248 |
| Cardiac Troponin T | rabbit | 1:200 (IF); 1:1000 (WB) | Abcam | ab45932 |
| α-Actinin EA53 | mouse | 1:400 (IF) | Sigma-Aldrich | A7811 |
| Ki-67 (D3B5) | rabbit | 1:400 (IF) | Cell signaling | #9129 |
| $\alpha/\beta$-Tubulin | rabbit | 1:500 (WB) | NEB | 21148 S |

### Immunocytochemistry of primary hepatocytes

Primary hepatocytes were fixed in 2% PFA for 15 min at room temperature. After three washes with PBS, nuclei were stained with Hoechst dye (1:5000 from a 10 mg/mL stock) for 30 min at RT. Images were acquired using either a CellInsight CX7 Platform (Thermo Scietific) or a Keyence BZ-9000E/X810 microscope.

For quantification, 6-well plates were scanned and the numbers of Venus- and Scar-let-positive cells were determined relative to the total cell number (Hoechst-positive cells) using the Thermo Scientific HCS Studio Cell Analysis or the Keyence BZ-X800 Analyzer software.

### Immunocytochemistry of primary tissues

Mice were euthanized and perfused with PBS followed by 2% PFA. Liver tissue was dissected and embedded in OCT medium (Medite), frozen, and sectioned at 5 µm using a CryoStar NX70 cryostat (Thermo Scientific). Sections were mounted on histological slides and stored at -80°C until further processing.

For staining, tissue sections were fixed on slides in 2% PFA for 10 min and permeabilized for 1 h in PBS containing 0.05% Tween-20 and 0.025% Triton X-100. Nuclei were stained with Hoechst dye (1:5000 dilution from a 10 mg/mL stock solution) in PBS for 30 min at room temperature. Sections were mounted using CC/Mount medium (Sigma), and images were acquired using a Keyence BZ-9000E microscope.

### Fluorescence intensity measurement

Control and REPLACE-treated mice were sacrificed and perfused with 2% PFA. Dissected tissues were collected and homogenized in Triton X-100 lysis buffer (20 mM HEPES, 300 mM NaCl, 0.2 mM EDTA 1.5 mM MgCl_2_, 1% Triton-X 100) supplemented with phosphatase and protease inhibitors (Roche). Tissue lysates corresponding to 100 μg total protein, diluted in PBS, were transferred in 96-well black microplates (Greiner BioOne), and fluorescence signals were measured using an Infinite M1000 Pro plate reader (Tecan).

### FACS analysis and sorting

Venus- and Scarlet-expressing primary hepatocytes obtained from in vivo experments were quantified and sorted using a BD Aria III flow cytometer (BD Biosciences). Briefly, primary hepatocytes were isolated from control and REPLACE-treated mice and collected in low-glucose DMEM (1 g/L glucose; Life Technologies). For quantification, at least 20,000 viable single cells were recorded per sample. For downstream genomic DNA extraction, 100,000–300,000 Scarlet-positive cells were sorted. Data were analyzed using FlowJo software (Tree Star, version 10.8.1).

### Western Blot

Protein lysates were obtained using RIPA buffer supplemented with phosphatase and protease inhibitors (Roche). Protein concentration was determined using the Pierce BCA Protein Assay Kit (Thermo Fisher) following the manufacturer’s instructions. 50 μg of protein were resolved on a 4-20% Mini-PROTEAN TGX Stain-Free Protein Gel (Bio-Rad) and transferred to a 0,45 μm nitrocellulose membrane (Fisher Scientific) using a Mini-PROTEAN Tetra Vertical Electrophoresis Cell (Bio-Rad). After blocking the membrane in 5% milk powder for one hour at room temperature, the primary antibody (Table 3) was applied overnight at 4°C. The respective horse radish peroxidase (HRP)-conjugated secondary antibody was applied by incubating for 1.5 hours at room temperature. For detection, membranes were incubated with Pierce ECL Western Blotting Substrate or SuperSignal West Femto Maximum Sensitivity Substrate (Thermo Fisher) and imaged in a ChemiDoc Imaging System (Bio-Rad).

### Statistics and reproducibility

Data was processed and statistical analysis was performed in python and GraphPad Prism (v10.2.2). Data are shown as means with standard deviations (SDs). Data distribution was assessed for normality using the Shapiro-Wilk test and Q-Q plot inspection. Sample sizes, p value, and the statistical tests used are described in the figure legends. A p value of less than 0.05 was considered statistically significant. No statistical method was used to predetermine sample size. No data were excluded from the analyses. The experiments were not randomized, and the investigators were not blinded to allocation during experiments and outcome assessment.

## Data and code availability

The main data are available in the main text or the supplemental information. Illumina sequencing data are available in the Sequence Read Archive under the BioProject accession number: PRJNA1366436. Additional data and/or additional files are available via corresponding authors upon reasonable request. The code sources for Lamin A/C fluorescence intensity quantification and PacBio sequencing analysis are available at:

https://gitlab.com/ida-mdc/brightness-measurements and

https://github.com/lebmih/hiros

## Acknowledgments

We thank the Advanced Light Microscopy and Flow Cytometry technology platforms at the Max Delbrück Center for Molecular Medicine in the Helmholtz Association, Berlin, Germany, for technical support and access to instruments. We gratefully acknowledge the excellent technical support of Andrea Leschke and Carola Guttwein from the Transgenic Core Facility, and of Carlon Genehr from the Technology Platform for Pluripotent Stem Cells at the Max Delbrück Center for Molecular Medicine in the Helmholtz Association, Berlin, Germany. This work was supported by the European Research Council (Horizon Pathfinder Challenges 01 - grant 101115295 – NaV1.5-CARED).

## Author contributions

T.D. and A.R. contributed equally to this work. T.D. and R.K. conceived the study and provided the conceptual framework. T.D. coordinated the project and oversaw experimental design and execution. T.D., A.R. and R.K. designed the study. T.D., A.R. and A.Z. performed the experiments with assistance from C.G., M.E. and S.W. T.D. and A.R. analyzed the data and wrote the original manuscript. M.D. and S.B. provided the iPSC-lines, established cardiac differentiation protocols and assisted with cardiac differentiation. M.L. analyzed the PacBio Long-Read sequencing data. E.B. wrote the scripts for quantification of Lamin A/C fluorescence intensity signals.

## Declaration of generative AI and AI-assisted Technologies in the writing process

During the preparation of this work, the authors used ChatGPT4 in order to refine sentence structure. After using this tool, the authors reviewed and edited the content as needed and take full responsibility for the content of the publication.

## Supplementary Figures

**Supplementary Figure 1.**
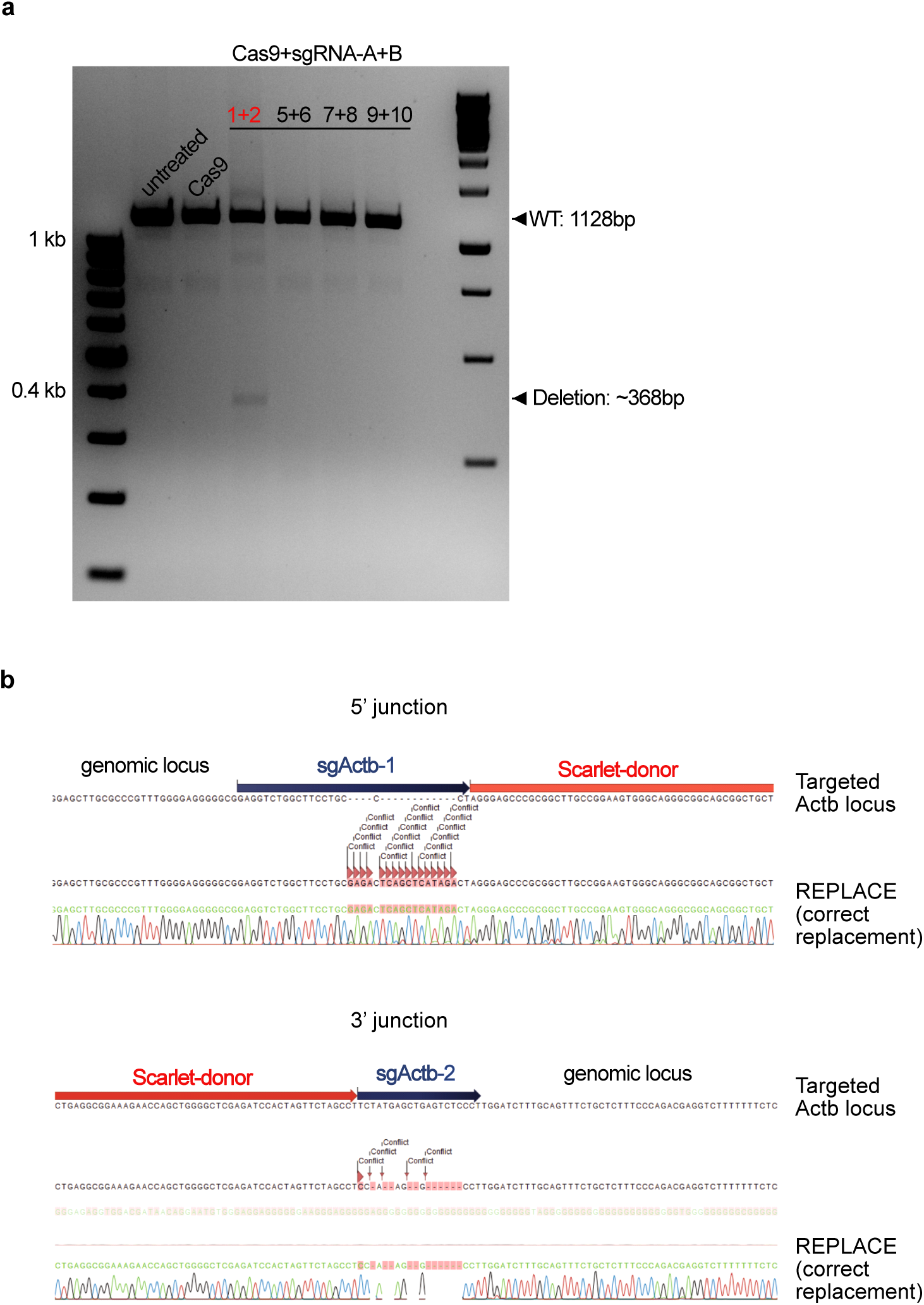
sgRNA selection and sequencing validation of ACTB-Scarlet REPLACE in mouse primary hepatocytes. **(a)** Screening of candidate sgRNA pairs targeting the mouse *Actb* locus. N2A cells were transfected with SpCas9 and the indicated sgRNA combinations, and the genomic interval spanning both cut sites was amplified by PCR. The unedited (WT) /one-cut amplicon (∼1128 bp) is present in all conditions, whereas successful dual cutting yields a shorter deletion product (∼368 bp). **(b)** Sanger sequencing confirma-tion of correct replacement junctions. PCR products spanning the 5′ and 3′ donor-genome junctions from primary hepatocytes treated with AAV-ACTB-donor and AAV-sgActb1+2 were sequenced and aligned to the expected edited allele.

**Supplementary Figure 2.**
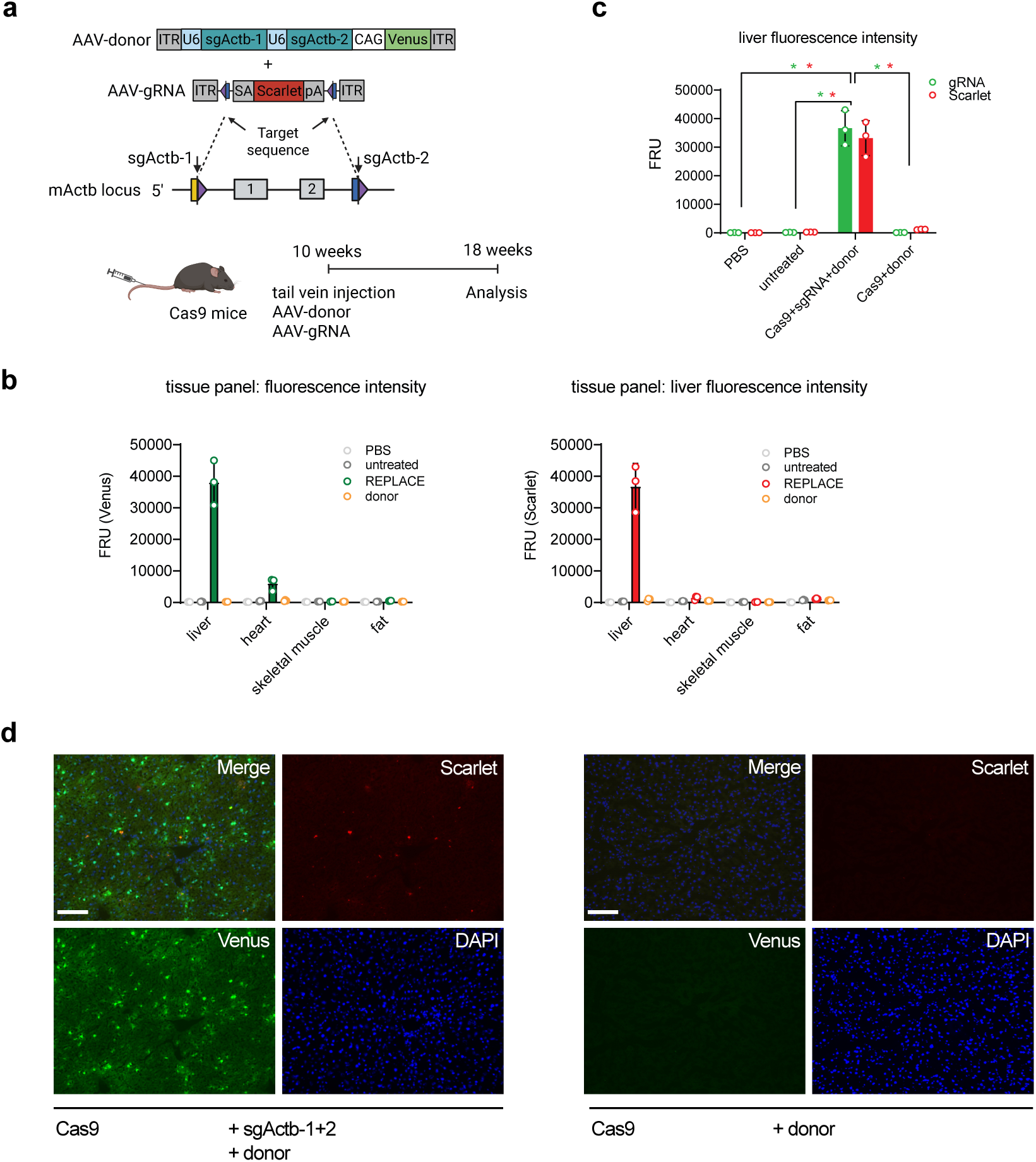
In vivo delivery and fluorescence expression after ACTB-Scarlet REPLACE in R26-Cas9 mice. **(a)** Experimental scheme for ACTB-Scarlet REPLACE in R26-Cas9 mice. Ten-week-old male R26-Cas9 animals were injected via tail vein with AAV-D/J vectors encoding the ACTB-donor and AAV-sgActb1+2 (Venus reporter). **(b)** Tissue panel (liver, heart, skeletal muscle, white adipose tissue) for Venus and Scarlet expression from mice 8 weeks after injection. Venus and Scarlet expression was examined by fluorometric measurement of Venus and Scarlet intensity (RFU). Data are expressed as mean ± SD (n = 3 mice per group). **(c)** Scarlet and Venus (gRNA) expression examined by fluorometric measurement (RFU) in liver from control mice (PBS injected, untreated, donor) and mice injected with AAV-ACTB-donor and AAV-sgActb1+2. Data are expressed as mean ± SD (n = 3 mice per group), statistical significance was determined by two-way ANOVA and Turkey’s multiple comparison test with * p ≤ 0.05. **(b)** Representative fluorescence images of liver cryosections from treated (AAV-ACTB-donor and AAV-sgActb1+2) and control animals (AAV-ACTB-donor-only). Scale bar, 100 µm.

**Supplementary Figure 3.**
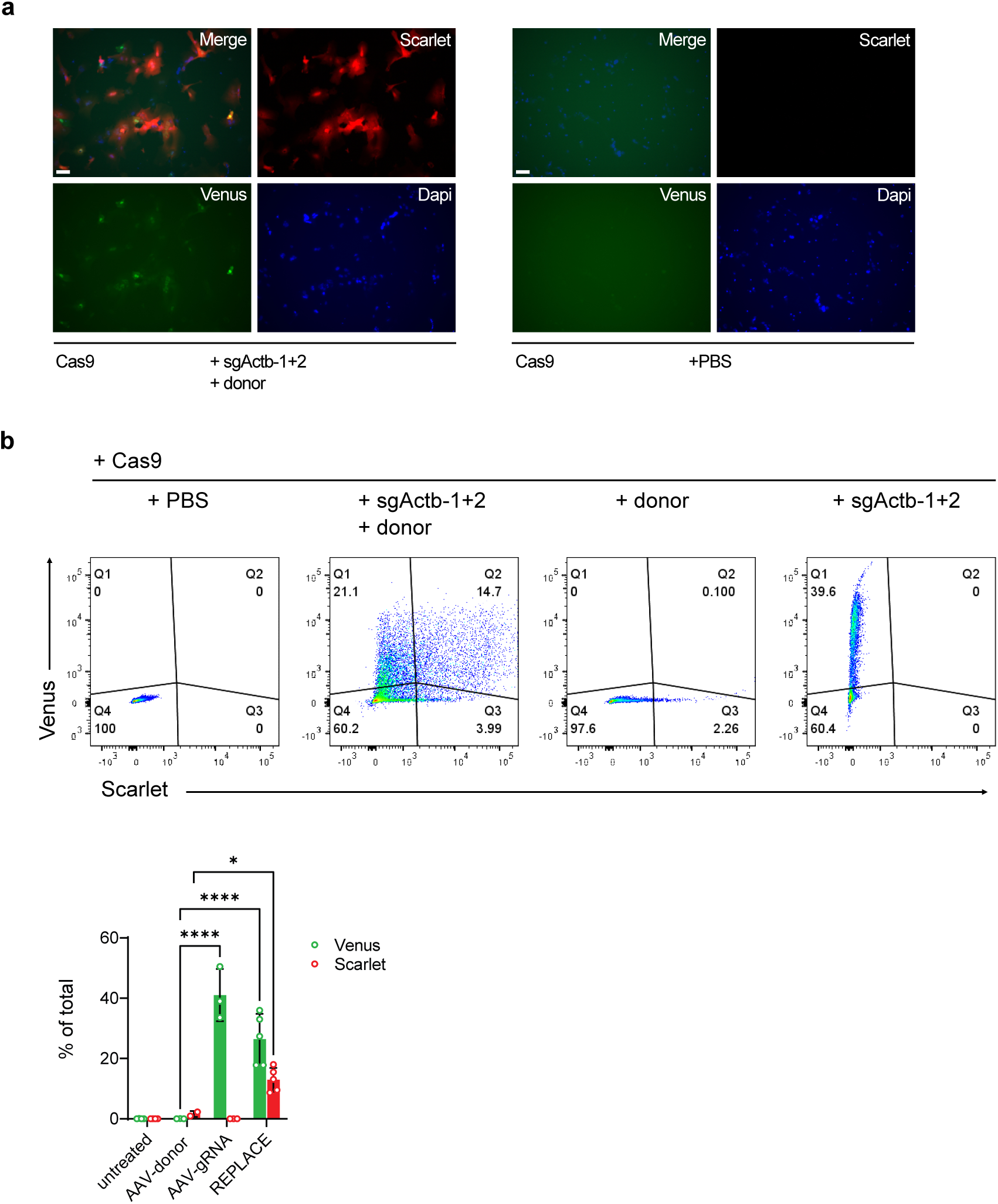
Quantification of in vivo ACTB-Scarlet REPLACE editing in R26-Cas9 mice. **(a)** Representative fluorescence images of primary hepatocytes isolated from control (PBS injected) and treated R26-Cas9 mice (AAV-ACTB-donor and AAV-sgActb1+2) 8 weeks after tail vein injection. Scale bar, 100µm. **(b)** Flow-cytometry analysis for quantifying transduction and editing outcomes in isolated hepatocytes from control and treated mice. The gates indicate the percentages of Scarlet+ and Venus+ cells. Data are expressed as mean ± SD (n = 3 mice per group), statistical significance was determined by two-way ANOVA and Dunnett’s multiple comparison test with * p ≤ 0.05, **** p ≤ 0.0001.

**Supplementary Figure 4.**
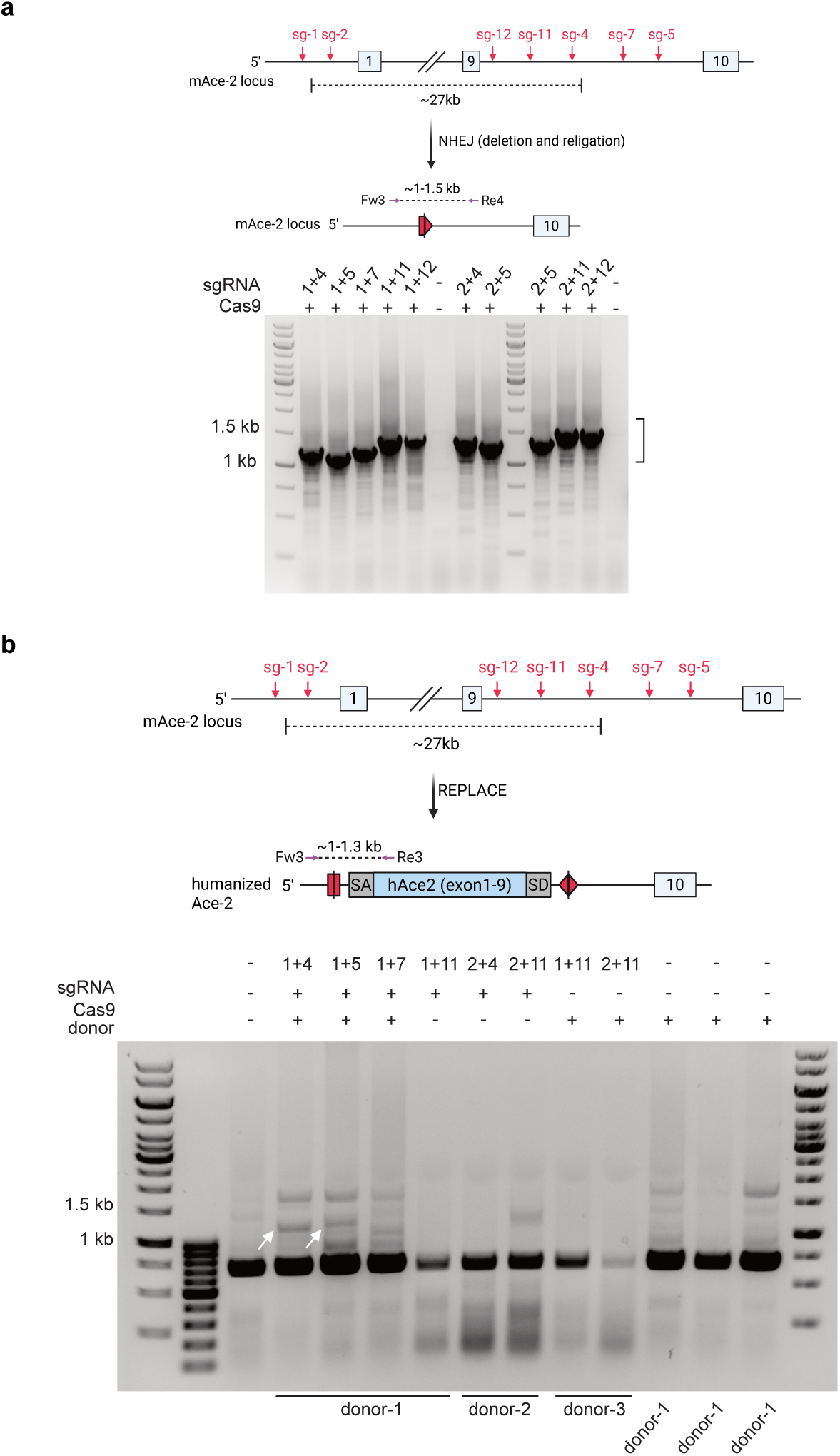
sgRNA screening and validation of hAce2-REPLACE strategy. **(a)** Screening of candidate sgRNAs pairs (red arrows) in N2A cells. Validation of sgRNA activity by PCR amplification of genomic interval spanning both cut sites using indicated primers (purple arrows: Fw3, Re4). Unedited locus exceeds 27 kb, no full-length amplicon was detected under standard PCR conditions. Productive dual cutting generated shorter deletion products of 1-1.5 kb as indicated by black bracket. **(b)** Validation of hAce2-REPLACE donor constructs in 19PP cells. Cas9/sgRNA RNPs and double stranded DNA donor generated by PCR were delivered by nucleofection. Donor-1 contains the inverted gRNA binding sites of sg-1 (sgAce2-1), donor-2 gRNA binding sites of sg-2 (sgAce2-2), and donor-3 contains the inverted gRNA binding sites of sg-11 (sgAce2-11). Junction PCR was performed using indicated primers (Fw3, Re3). White arrows indicate correct integration.

**Supplementary Figure 5.**
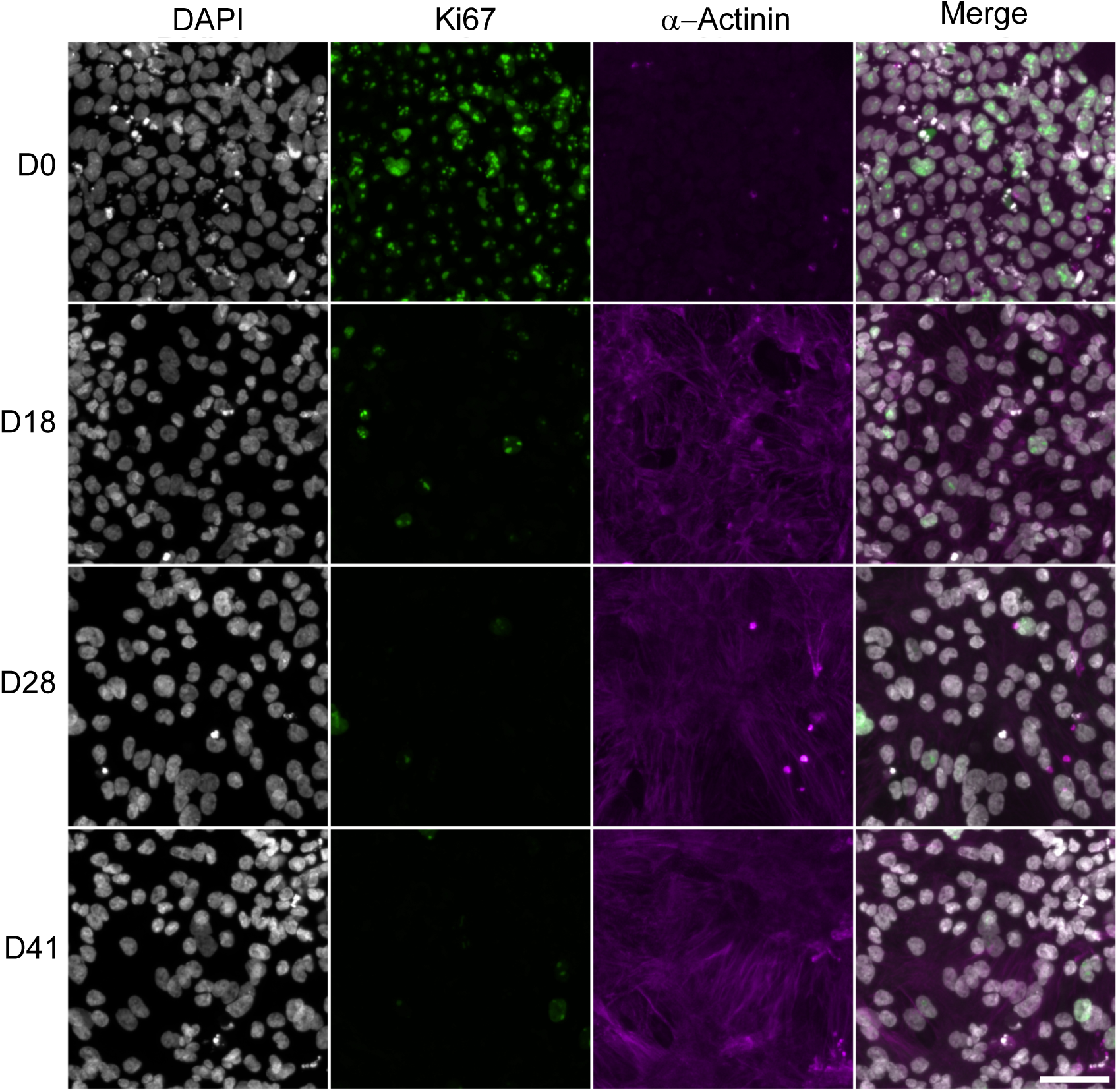
Immunocytochemistry staining of proliferation and cardiac marker in mutant hi-CMs. Representative images of homozygous LMNAk117fs-hiPSCs (day 0) and hi-CMs at day 18, 28 and 41. Cells were stained for DAPI (grey), Ki67 (green) and α-Actinin (magenta). Scale bar, 50 μm.

**Supplementary Figure 6.**
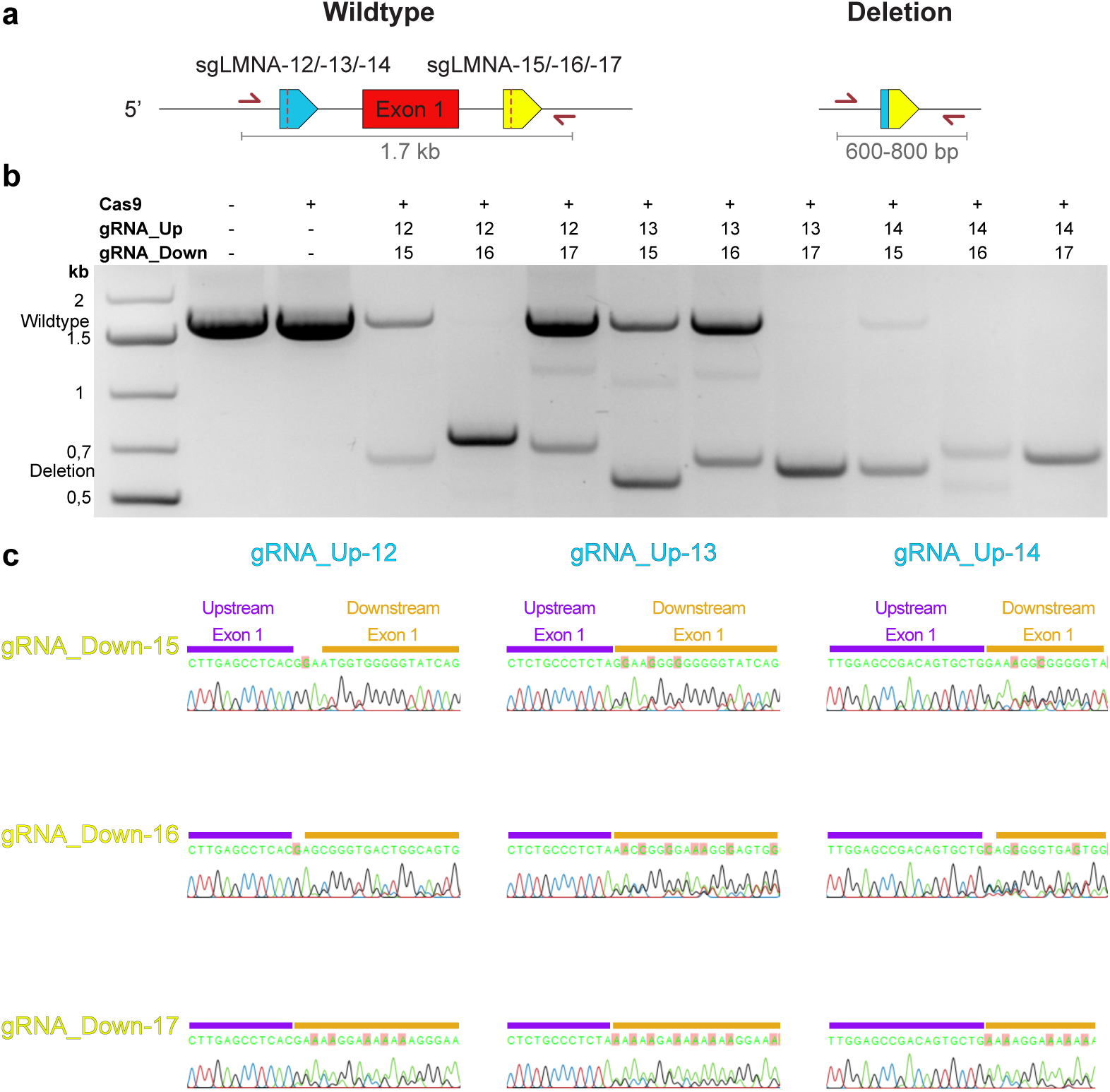
sgRNA screening for LMNA-REPLACE strategy. **(a)** Schematic illustration of the genotyping PCR used to identify activity of up- and downstream-sgRNA (gRNA Up/ Down). Amplification with primers (red half arrows) binding outside the LMNA-sgRNA target sites produces large 1.7 kb amplicons from unedited wildtype alleles (left). Robust activity of both LMNA-sgRNAs leads to the deletion of exon 1, which gives rise to shorter PCR amplicons between 600 and 800 bp, depending on the sgRNA pair (right). **(b)** Agarose gel of PCR amplicons as described in A. **(c)** Sanger sequencing of the re-ligation sites in the short PCR amplicons.

